# Perinatal Programming of Mucosal Stromal Cell Identity by the Lymphotoxin Pathway Regulates Mucosal Immune Responses in the Adult

**DOI:** 10.1101/557652

**Authors:** Conglei Li, Evelyn Lam, Christian Perez-Shibayama, Dennis Lee, Lesley A. Ward, Albert Nguyen, Musaddeque Ahmed, Kirill V. Korneev, Olga Rojas, Tian Sun, Housheng Hansen He, Alberto Martin, Burkhard Ludewig, Jennifer L. Gommerman

**Author notes:** Corresponding author: Jennifer L. Gommerman, Ph.D., Department of Immunology, University of Toronto, Medical Sciences Building, 1 King’s College Circle, Room 7233, Toronto, Ontario, Canada M5S 1A8.

## Abstract

Redundant mechanisms support IgA responses to intestinal antigens. These include multiple priming sites (mesenteric lymph nodes (MLN), Peyer’s patches and isolated lymphoid follicles) and various cytokines that promote class switch to IgA, even in the absence of T cells. In spite of these back-up mechanisms, vaccination against enteric pathogens such as Rotavirus has limited success in some populations. Genetic and environmental signals experienced during early life are known to influence mucosal immunity, yet the mechanisms for how these exposures operate remain unclear. Here we used Rotavirus infection to follow antigen-specific IgA responses through time and in different gut compartments. Using genetic and pharmacological approaches, we tested the role of a pathway known to support IgA responses (Lymphotoxin – LT) at different developmental stages. We found that LT-beta receptor (LTβR) signalling *in utero* programs intestinal IgA responses in adulthood by affecting antibody class switch recombination to IgA and subsequent generation of IgA antibody-secreting cells within an intact MLN. In addition, *in utero* LTβR signalling dictates the phenotype and function of MLN stromal cells in order to support IgA responses in the adult. Collectively, our studies uncover new mechanistic insights into how *in utero* LTβR signalling impacts mucosal immune responses during adulthood.

**One Sentence Summary:** Early life LTβR signalling is critical for programming the mesenteric lymph node stromal cell environment, impacting both antibody isotype switching to IgA and the differentiation of IgA^+^ antibody secreting cells.

*Graphic Abstract:* 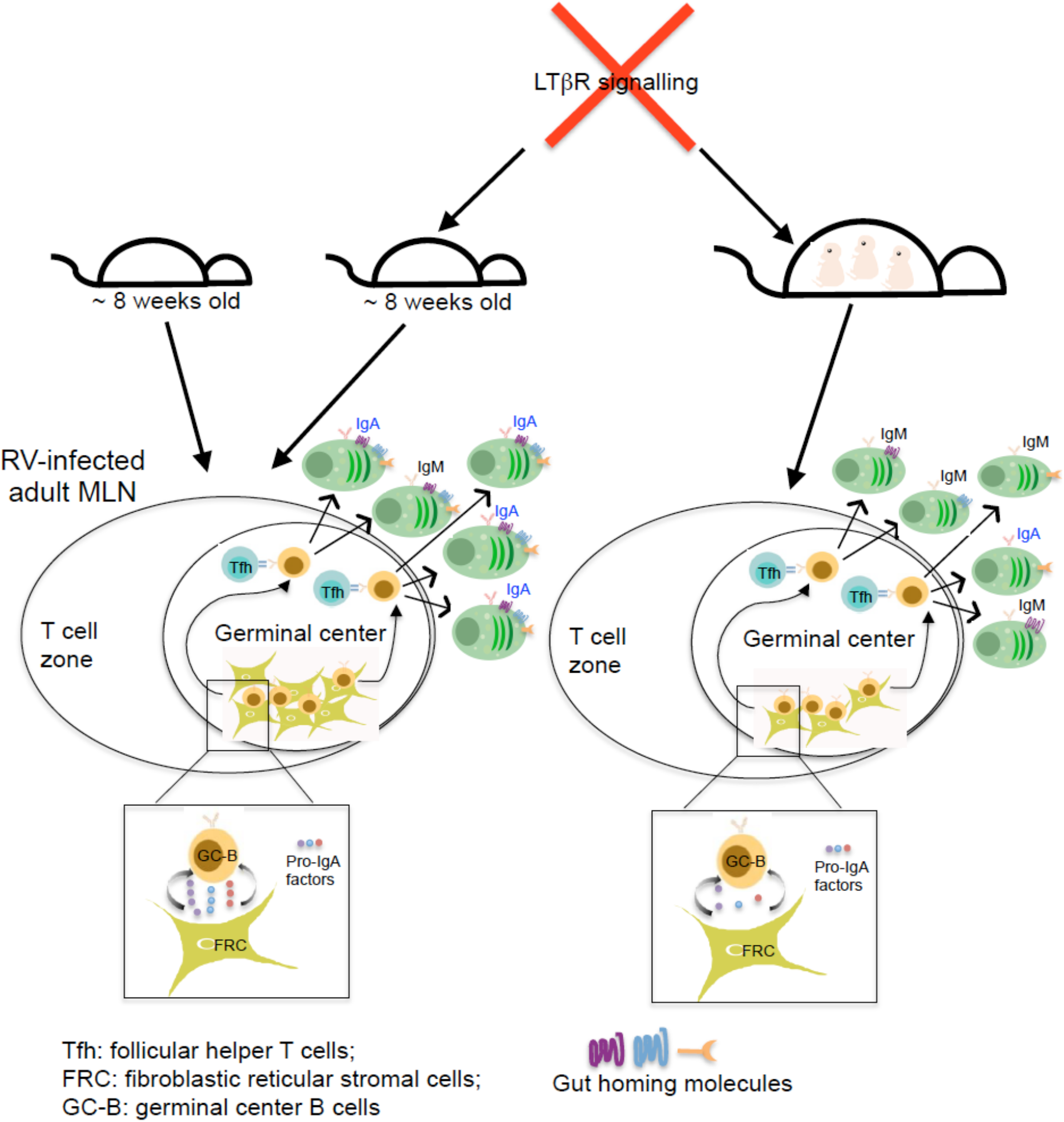

## INTRODUCTION

Lymphotoxin-beta Receptor (LTβR) signalling is required for the development of secondary lymphoid organs *in utero (1-3*), and for the maintenance of lymphoid tissue architecture during adulthood (*4, 5*). The membrane-bound form of the LTβR ligand (LTα_1_β_2_) is expressed by activated T and B lymphocytes as well as RORγt^+^ innate lymphoid cells, while the corresponding LTβR is expressed by dendritic cells (DC), macrophages, and radio-resistant stromal cells (*4, 6, 7*). LTα^-/-^ and LTβR^-/-^ mice lack PP and all LNs (*1, 8, 9*), whereas LTβ^-/-^ mice lack PP and most LNs, but retain some mucosal-associated LNs including MLN (*2, 10*). While serum IgM and IgG levels are relatively normal in LT-deficient mice, both serum and intestinal IgA levels are severely reduced (*9-11*). The selective reduction in IgA is not due to a lack of secondary lymphoid organs in LT-deficient mice (*11, 12*), raising the key question of what is the mechanism of LTβR-dependent control of IgA responses.

IgA is the most abundant antibody isotype in the body and more than 80% of human plasma cells produce IgA (*13-15*). IgA production at mucosal sites is a major means for containing the intestinal microbiota and preventing the colonization of infectious agents, such as pathogenic viruses (*14-16).* Intestinal IgA can be generated via both T cell-dependent and T-independent mechanisms (*13, 17, 18*). T-dependent IgA responses are initiated within gut-associated lymphoid tissue (mesenteric lymph nodes – MLN, and Peyer’s patches – PP) via the formation of germinal centers (GC) which support interactions between B cells and follicular helper T cells (Tfh) (*13*). Post-germinal center IgA^+^ antibody secreting cells (ASC) egress through efferent lymphatics and subsequently enter the blood via the thoracic duct. Because of their expression of homing chemokine receptors and adhesion molecules such as CCR9, CCR10 and α4β7 integrin (*19, 20*), ASC primed in the MLN or PPs ultimately circulate back to the intestinal lamina propria effector site where they secrete IgA that is transported into the gut lumen.

Recent studies have reported that DC-intrinsic LTβR signalling contributes to local IgA production within PP in response to the microbiota (*21*), whereas other studies point to the role of LTβR signalling in the radio-resistant compartment (*11, 12).* Whether intestinal IgA responses are initiated by T-dependent *versus* T-independent mechanisms, and in what location these responses are intitiated (MLN, PP, LP), are key variables that likely influence how LT and related pathways impact mucosal IgA responses (*22-24*). Due to these variables, model systems that measure *de novo* IgA responses to foreign enteric pathogens are a useful strategy for answering questions about how IgA responses are generated in the host. When and how the LT pathway shapes the IgA response to a foreign mucosal pathogen has not been fully addressed.

One very important and emerging parameter that influences mucosal immune responses is exposure to perinatal signals which comprise inputs from both environmental and genetic factors (*25, 26*). How such signals play a role in influencing mucosal IgA responses is unclear. Given that LTβR plays a critical role in supporting lymphoid tissue development during fetal life (*27*), we hypothesized that *in utero* LTβR signals are likewise important for programming the identity of key support cells that influence immune responses within intact lymphoid tissues in the adult animal. To test this, we took advantage of genetic and pharmacological approaches that inhibit LTβR signalling yet spare the development of MLN. We combined this approach with an infection model (Rotavirus –s RV) that we show is dependent on B cell priming within the MLN. Using this strategy, we found that LTβR signalling during fetal life, but not during adulthood, is critically required to generate a *de novo* antigen-specific IgA response in the gut via programming the MLN stromal cell environment. Collectively, our studies provide new insights into how early life signalling events impact the identity of gut-resident stromal cells as well as responses to foreign pathogens during adulthood.

## RESULTS

### Mice that lack LTβ in early life exhibit impaired RV-specific IgA responses during adulthood

IgA levels in LT-deficient mice, which lack PP and LN, are profoundly suppressed (*9-11*), and introduction of LT-sufficient bone marrow (BM) into LT-deficient adult mice has been shown to rescue polyclonal IgA responses via signalling through LTβR in the gut (*11*). While these results imply that at least some IgA responses do not require early life LTβR signalling, it is unclear if the lack of dependency on early life LTβR signals would also be true in the context of IgA responses to mucosal pathogens. We selected Rotavirus (RV) as a model pathogen for several reasons: In mice, RV is highly trophic for small intestinal epithelial cells and induces a predominant IgA response that peaks in the first 9-13 days (d) post-infection (*28, 29*). Although RV infection in neonates can cause mild small intestinal pathology and diarrhea, RV infection in adults is asymptomatic (*30*). Thus, RV infection is an appealing model for studying a *de novo* IgA response in the adult gut without accompanying pathology.

To test the role of early life LTβR signalling on polyclonal and RV-specific IgA responses, we created BM chimeric mice whereby WT or LTβ^-/-^ BM was introduced into WT or LTβ^-/-^ recipient mice, followed by co-housing (*31*) (**Fig. 1a, and Supplementary Fig. 1a**). We then examined levels of polyclonal IgA in the feces (presumably directed at commensal microbiota) *versus* IgA responses to RV. As expected, LTβ^-/-^ mice that received LTβ^-/-^ BM had very low levels of fecal IgA. Moreover, consistent with the results of Kang et al (*11*), we observed comparable levels of total fecal IgA in WT➔LTβ^-/-^ and WT➔WT chimeric mice (**Supplementary Fig. 1b**), confirming that restoration of LTβR signalling in adult mice (by introduction of LTβ^+/+^ BM) has the capacity to rescue IgA levels in the gut. We next measured anti-RV IgA levels in mice that lack LTβR signalling in early life. Adult WT mice housed in our vivarium exhibit fecal anti-RV IgA levels at approximately d7 post-infection concomitant with RV antigen clearance (**Supplementary Fig. 1c-d**). As there is no anti-RV antibody standard available for this ELISA assay, all the fecal samples were tested in one ELISA plate, and results were confirmed using ELISPOT (see below). Interestingly, in contrast to the observed normalization of polyclonal fecal IgA levels upon transplantation of WT BM into LTβ^-/-^ mice (**Supplementary Fig. 1b**), WT➔LTβ^-/-^ chimeras exhibited a significant reduction in their ability to mount an anti-RV IgA response that persisted from d8 to d49, when compared to WT➔WT controls, and the low levels of anti-RV IgA in WT➔LTβ^-/-^ chimeras were comparable to LTβ^-/-^ ➔LTβ^-/-^ chimeras (**Fig. 1b**). In addition, we employed an RV-specific ELISPOT to assay the frequency of IgA^+^ RV-ASC in the SILP at d8 and d49 post-RV infection, respectively. In comparison to WT➔WT controls, the accumulation of IgA^+^ RV-ASC in the SILP was also significantly reduced in both WT➔LTβ^-/-^ and LTβ^-/-^➔LTβ^-/-^ chimeras at d8 and d49 postinfection, respectively (**Fig. 1c**). Taken together, these results demonstrate that early life LTβR signalling is essential for anti-RV IgA responses during adulthood.

**Fig. 1.**
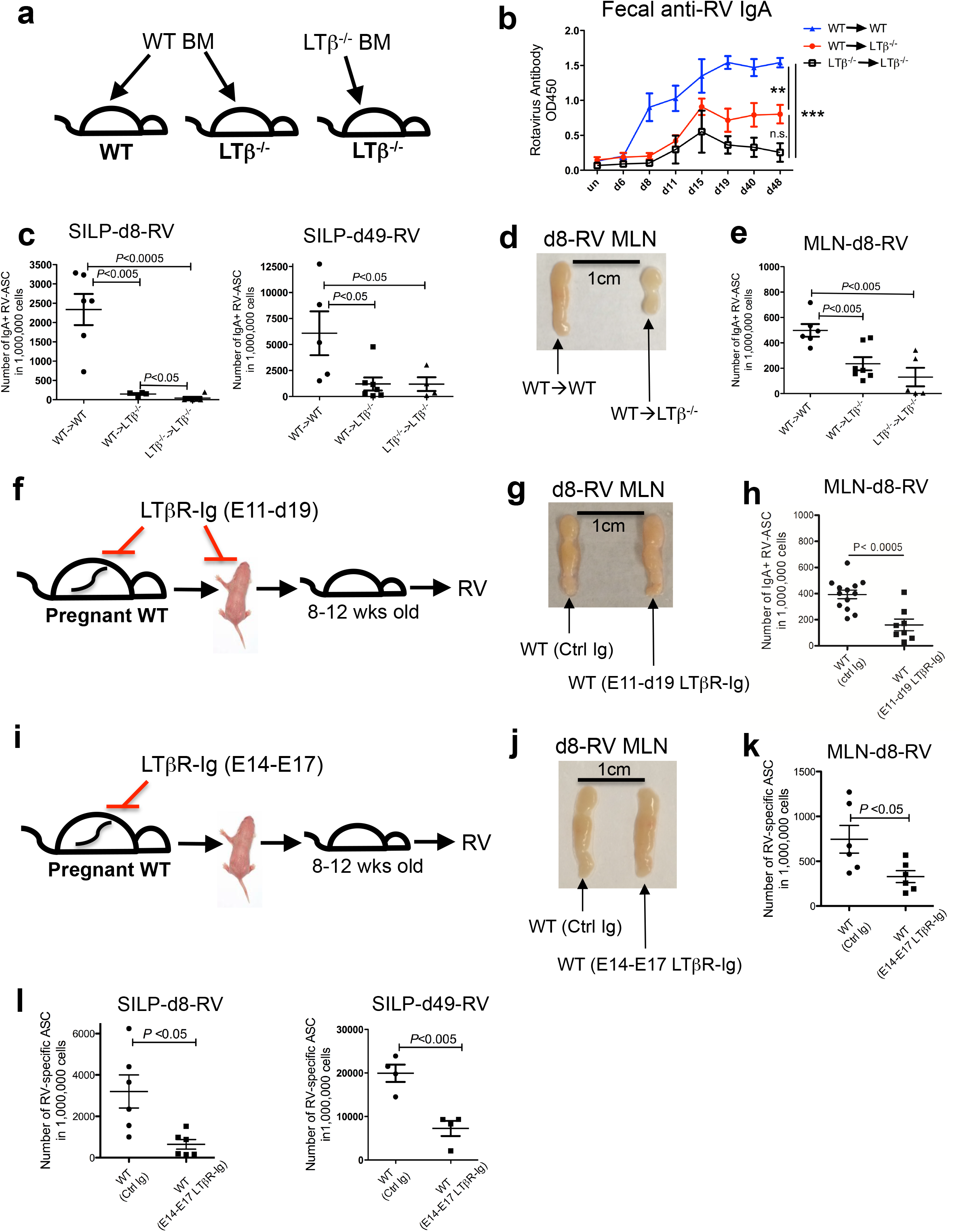
LTβR signalling during fetal life in mice with intact MLN is critically required for the generation of a RV-specific IgA response. **(a)** Diagram depicting experimental setup. **(b)** ELISA analysis of fecal anti-RV IgA levels in co-caged BM chimeric mice. Data represent two independent experiments each with four mice per time point and were analyzed by two-way ANOVA (* P<0.05; ** P<0.01; ***P<0.001; n.s. = not significant). All the samples were tested in one ELISA plate. **(c)** Enumeration of IgA^+^ RV-ASC in the SILP of BM chimeric mice at d8 (left panel) and d49 (right panel) post-infection. Data represent at least two independent experiments each with four to six mice per group. (d) Representative picture of MLN from WT➔WT *versus* WT➔LTβ^-/-^ chimeric mice at d8 post-infection. **(e)** Enumeration of IgA^+^ RV-ASC in the MLN of chimeric mice at d8 post-infection. Data represent at least three independent experiments each with four to six mice per group. **(f)** Depiction of experimental setup for inhibiting LTβR signalling during both fetal and neonatal periods (E11-d19). **(g)** Representative picture of MLN from E11-d19 LTβR-Ig or control Ig (Ctrl Ig) treated WT mice at d8 postinfection. **(h)** Analysis of IgA^+^ RV-ASC in the MLN of E11-d19 LTβR-Ig or control Ig treated WT mice (n= 8-12 mice per group; data were pooled from 3 independent experiments). **(i)** Depiction of experimental setup for *in utero* (E14/E17) inhibition of LTβR signalling in WT mice. **(j)** Representative picture of MLN from E14/E17 LTβR-Ig or control Ig (Ctrl Ig) treated WT mice at d8 post-infection. **(k)** Enumeration of IgA^+^ RV-ASC in the MLN at d8 postinfection of WT mice that received LTβR-Ig or control Ig at E14/E17. **(l)** Enumeration of IgA^+^ RV-ASC in the SILP at d8 (left panel) and d49 (right panel) post-infection of WT mice that received LTβR-Ig or control Ig at E14/E17. Data represent at least three independent experiments each with four to five mice per group.

### LTβR signalling is critically required for the generation of an anti-RV IgA response in the adult MLN

We next asked whether the defect in production of RV-specific IgA observed in the SILP and the fecal pellet of WT➔LTβ^-/-^ mice was due to a problem at the effector site (SILP) or due to a defect in the priming site. To ascertain which possibility applied, we first determined the location of B cell priming against RV in our mice. Since activation-induced cytidine deaminase (AID) is triggered by BCR activation (*32*), we used an AID-GFP reporter mouse to identify the location of B cell priming against RV (*33, 34*). Because fecal anti-RV IgA was induced at approximately d7 post-infection in WT mice (**Supplementary Fig. 1c**), we assayed the expression of AID-GFP in B cells within this time-frame. AID-GFP transgenic mice that were infected with RV showed evidence of AID-GFP expression mainly in the MLN rather than the PP or the SILP (**Supplementary Fig. 2a-b**). In addition, using our RV-specific ELISPOT readout we found that the MLN contained approximately 1000-fold more IgA^+^ RV-ASC compared to the PP at d8 postinfection, and 4-fold greater compared to the SILP at this time point (**Supplementary Fig. 2c**). Because there was limited AID-GFP induction in the SILP, presumably the IgA^+^ RV-ASC in the SILP represent recent immigrants from the MLN inductive site. Taken together, these data demonstrate that the MLN is the main inductive site for the initiation of an IgA class switched B cell response against RV.

We next determined if the defect in anti-RV IgA responses in WT➔LTβ^-/-^ mice was due to the generation of RV-specific IgA ASC in this location. We were able to assess this because WT➔LTβ^-/-^ mice retain a small MLN (**Fig. 1d**) (*10*). Accordingly, we assayed the number of IgA^+^ RV-ASC in the MLN of WT➔LTβ^-/-^ and LTβ^-/-^➔LTβ^-/-^ BM chimeras, comparing to WT➔WT controls. Interestingly, compared to WT➔WT controls, the number of RV-IgA^+^ ASC in the MLN was significantly reduced in WT➔LTβ^-/-^ chimeric mice, and to a similar extent as LTβ^-/-^➔LTβ^-/-^ chimeric mice (**Fig. 1e**). This suggests that the MLN environment in WT➔LTβ^-/-^ chimeric mice is incapable of supporting an RV-specific IgA response.

Thus far our results indicate that LTβ expression during the first 6 weeks of life is required for the induction of an anti-RV IgA response in an adult that has had LTβR signalling restored via WT BM transplantation. To more accurately narrow down the relevant period of LTβR signalling that is required for generating an anti-RV IgA response in the MLN, we treated pregnant WT dams and their neonatal offspring with LTβR-Ig, starting from embryonic d11 to postnatal d19 (E11-d19) (**Fig. 1f**) (*35*). We observed that, while most males develop MLN (6/7; compared to 7/7 for control Ig group), the majority of female littermates failed to develop MLN when treated with LTβR-Ig from E11-d19 (2/6 females have MLNs; compared to 6/6 for control Ig group, *P* < 0.05). Of those mice receiving E11-d19 LTβR-Ig treatment that successfully developed MLN, the size of their MLN was comparable to control Ig-treated mice (**Fig. 1g**). Nevertheless, in spite of a grossly normal appearing MLN, the generation of IgA^+^ RV-ASC in MLN was significantly impaired at d8 post-RV infection in E11-d19 LTβR-Ig treated mice compared to the control Ig treatment group (**Fig. 1h**).

To further narrow down the relevant LTβR-dependent developmental period that is required for an optimal IgA response to RV during adulthood, we treated pregnant WT mice with LTβR-Ig at E14 and E17 *(in utero* treatment) (**Fig. 1i**). For more than 50 mice examined, all E14/E17 LTβR-Ig treated mice successfully developed MLN, and MLN sizes were comparable to those derived from the control Ig treatment group (**Fig. 1j and Supplementary Fig. 2d**). Consistent with previous findings (*11*), we observed comparable anti-commensal polyclonal IgA responses in E14/E17 LTβR-Ig *versus* control Ig treatment groups (data not shown). However, we found that the frequency of antigen-specific IgA^+^ RV-ASC at d8-post RV infection was significantly reduced in the MLN of mice that received LTβR-Ig *in utero* compared to those that received control Ig (**Fig. 1k**). Consistent with this observation, the accumulation of IgA^+^ RV-ASC in the SILP was also significantly diminished at d8 and d49 post-RV infection in E14/E17 LTβR-Ig treated WT mice (**Fig. 1l**). In contrast, inhibition of LTαβ/LTβR signalling in adulthood, either by introduction of LTβ^-/-^ BM into adult WT mice or by treatment of adult mice with LTβR-Ig, had no impact on the anti-RV IgA response (**Supplementary Fig. 3**). In summary, our results show that *in utero* LTβR signalling from E14 to birth is required to support an optimal anti-RV IgA response during adulthood. Moreover, LTβR signalling from E11 (but not E14) is partially required for the development of MLN in female mice.

### *In utero* LTβR signalling is not required for the initiation of RV-induced GC responses in the adult MLN

There are many checkpoints that lead to the production of IgA-switched ASC during a mucosal immune response including the initiation of a GC response, upregulation of AID, class switch of B cells and their differentiation into PC that can home appropriately to the SILP. We therefore wished to understand how *in utero* LTβR signalling impacts the different steps that ultimately generate gut-homing anti-RV IgA producing plasma cells (PC). Accordingly, we first examined the frequency of discrete GC B cell subsets in the MLN as a percentage of CD19+ B cells (*21*) (**Supplementary Fig. 4a-b**). Compared to uninfected controls, the percentage of GC B cells was significantly increased in the MLN of WT➔LTβ^-/-^ chimeras at d8 post-RV infection (**Supplementary Fig. 4c**), and WT➔LTβ^-/-^ chimeras did not exhibit a significant defect in the percentage of GC or pre-GC/memory B cells in the MLN when compared to WT➔WT control chimeras (**Fig. 2a-b**). A comparable frequency of GC or pre-GC/memory B cells was also observed in mice treated *in utero* (E14/E17) with LTβR-Ig *versus* control Ig (**Fig. 2c**). Consistent with the notion that the defect in the MLN of WT➔LTβ^-/-^ mice is not due to an impaired GC response, RV infection induced a significant accumulation of Tfh in the MLN at d8 postinfection, and no differences in the accumulation of Tfh in the MLN were observed in WT➔LTβ^-/-^ *versus* control WT➔WT mice (**Supplementary Fig. 4d-e**). These data demonstrate that *in utero* LTβR signalling is dispensable for the initiation of a GC response in the MLN.

**Fig. 2.**
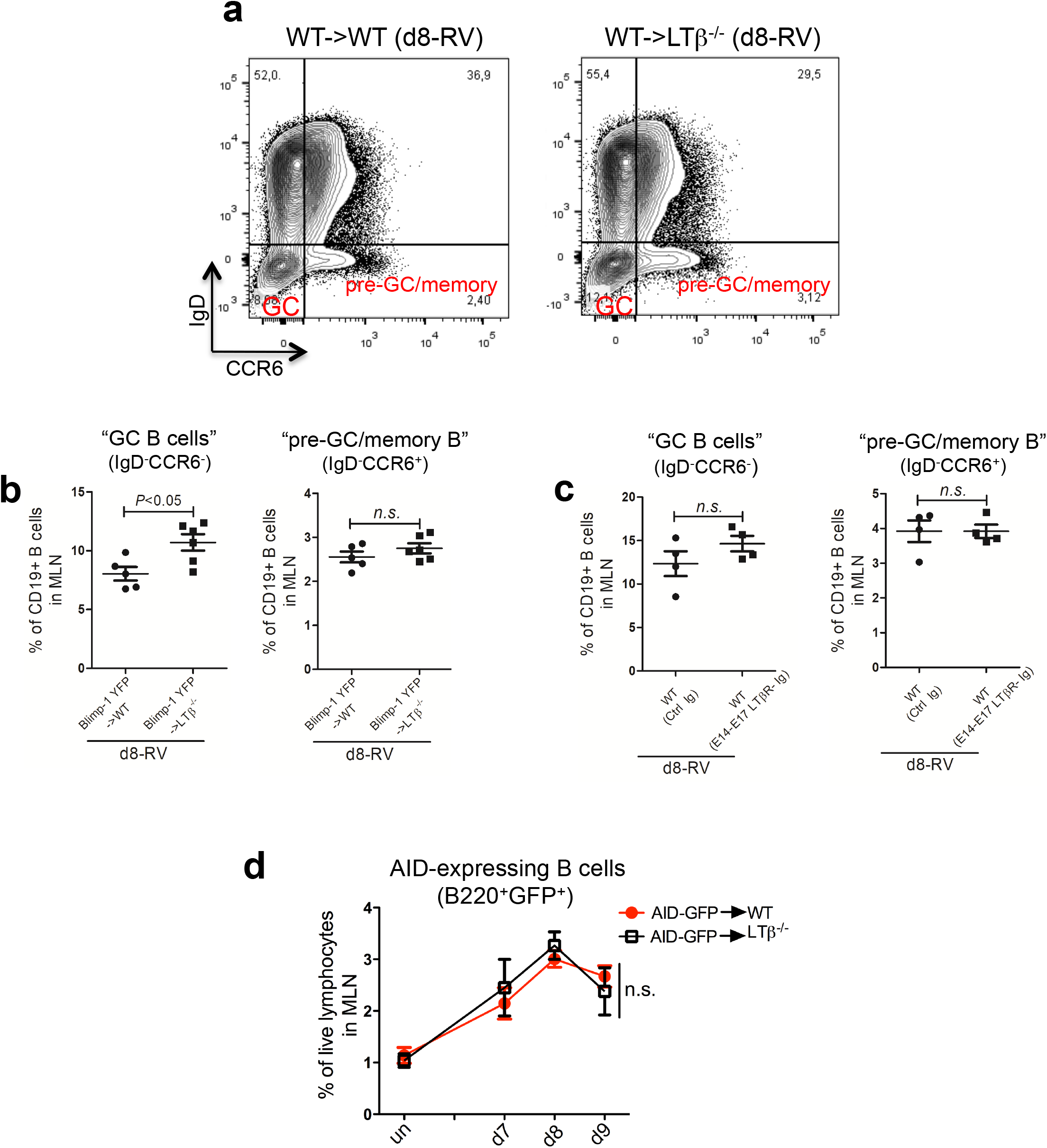
Early life LTβR signalling is not required for the initiation of RV-induced GC response in the adult MLN. **(a)** Representative FACS plots of germinal center (GC) B cells in the MLN of WT➔WT *versus* WT➔LTβ^-/-^ chimeras at d8 post-infection. **(b)** Frequency of GC B cells and pre-GC/memory B cells in MLN CD19+ cells were compared between WT➔WT *versus* WT➔LTβ^-/-^ chimeras. **(c)** Frequency of GC B cells and pre-GC/memory B cells in MLN CD19+ cells were compared between *in utero* (E14/17) LTβR-Ig and control Ig treated WT mice. **(d)** Frequency of AID-GFP+ B cells during the first 9 days post-infection in the MLN of the indicated BM chimeras (un: uninfected) (n=3 mice for uninfected group; n= 4-6 mice for RV infected group per time point). % = Frequency.

Transcription of AID in activated B cells is essential for antibody class switch recombination (CSR) from IgM to IgA in response to RV infection (*32*). To test whether the MLN of mice that lack LTβR signalling can support B-cell intrinsic expression of AID, we assayed the kinetics of AID-GFP expression in MLN B cells derived from AID-GFP➔WT *versus* AID-GFP➔LTβ^-/-^ chimeric mice. In spite of the lack of LTαβ/LTβR signalling during early life, we observed that the frequency of AID-expressing B cells in response to RV infection in the MLN of AID-GFP➔LTβ^-/-^ mice was comparable to that of AID-GFP➔WT mice (**Fig. 2d**). Thus, early life LTβR signalling is not required for the expression of AID in MLN-intrinsic B cells in response to RV infection.

In summary, although there is a profound reduction in IgA^+^ RV-specific ASC in the MLN of mice that lack LTβR signalling *in utero*, this cannot be explained by a defective GC response. Moreover, in spite of the observed reduction in the size of MLN in WT➔LTβ^-/-^ chimeric mice (**Fig. 1d**), the LTβ^-/-^ MLN (as well as MLN from mice treated with LTβR-Ig *in utero*) is capable of supporting at least the initial stages of a GC response, including the accumulation of Tfh, pre-GC B cells and GC B cells and the B cell-intrinsic induction of AID expression in response to RV infection.

### *In utero* LTβR signalling is required for IgA class switch and the accumulation of gut-homing plasma cells in response to RV infection in the adult mouse

Although AID expression is required for a class switched response to IgA, it is not sufficient. Thus, in spite of the normal GC response and AID induction in the LTβ^-/-^ MLN, it is possible that CSR to IgA in mice that lack early life LTβR signalling may still be impaired. We therefore assessed class switch to IgA in WT➔LTβ^-/-^ *versus* LTβ^-/-^➔LTβ^-/-^ chimeras as well as in mice treated *in utero* (E14/E17) with LTβR-Ig. Interestingly, compared to WT➔WT controls, we observed a marked increase in the frequency of IgM+ RV-ASC and a corresponding reduction in the frequency of IgA^+^ RV-ASC in the MLN of WT➔LTβ^-/-^ chimeric mice that translated into an overall reduction in the ratio of IgA^+^ *versus* IgM+ RV ASC, indicating a severe IgA CSR defect in MLN B cells from WT➔LTβ^-/-^ chimeric mice (**Fig. 3a-b**). In addition, compared to control Ig treatment, the IgA CSR defect was also observed when mice were treated with LTβR-Ig from E11-d19 as well as during the more narrow E14/E17 treatment window (**Fig. 3c-d**). Since AID expression was normal in AID-GFP ➔LTβ^-/-^ chimeric mice, we reasoned that the expression of the μ and α switch regions located upstream of the μ and α constant regions must be impacted by the absence of early life LTβR signalling in order to account for the observed defect in IgA CSR. To test this, we sorted out AID-GFP+B220+ cells from the MLN of AID-GFP ➔WT *versus* AID-GFP➔LTβ^-/-^ chimeric mice and evaluated the levels of Iμ-Cμ and Iα-Cα germline transcripts at d8 post-RV infection. Consistent with the observation of defective class switch in mice that lack early life LTβR signalling, we found that the level of Iα-Cα germline transcripts in MLN B cells from AID-GFP ➔LTβ^-/-^ chimeras was significantly reduced and accompanied by a trend towards reduced levels of Iμ-Cμ germline transcripts compared to AID-GFP➔WT controls (**Fig. 3e**). Collectively these data demonstrate that LTβR signalling from E14 to birth is required for the induction of Iα-Cα and Iμ-Cμ transcripts in AID+ B cells and subsequent IgA CSR in the adult MLN in response to RV infection.

**Fig. 3.**
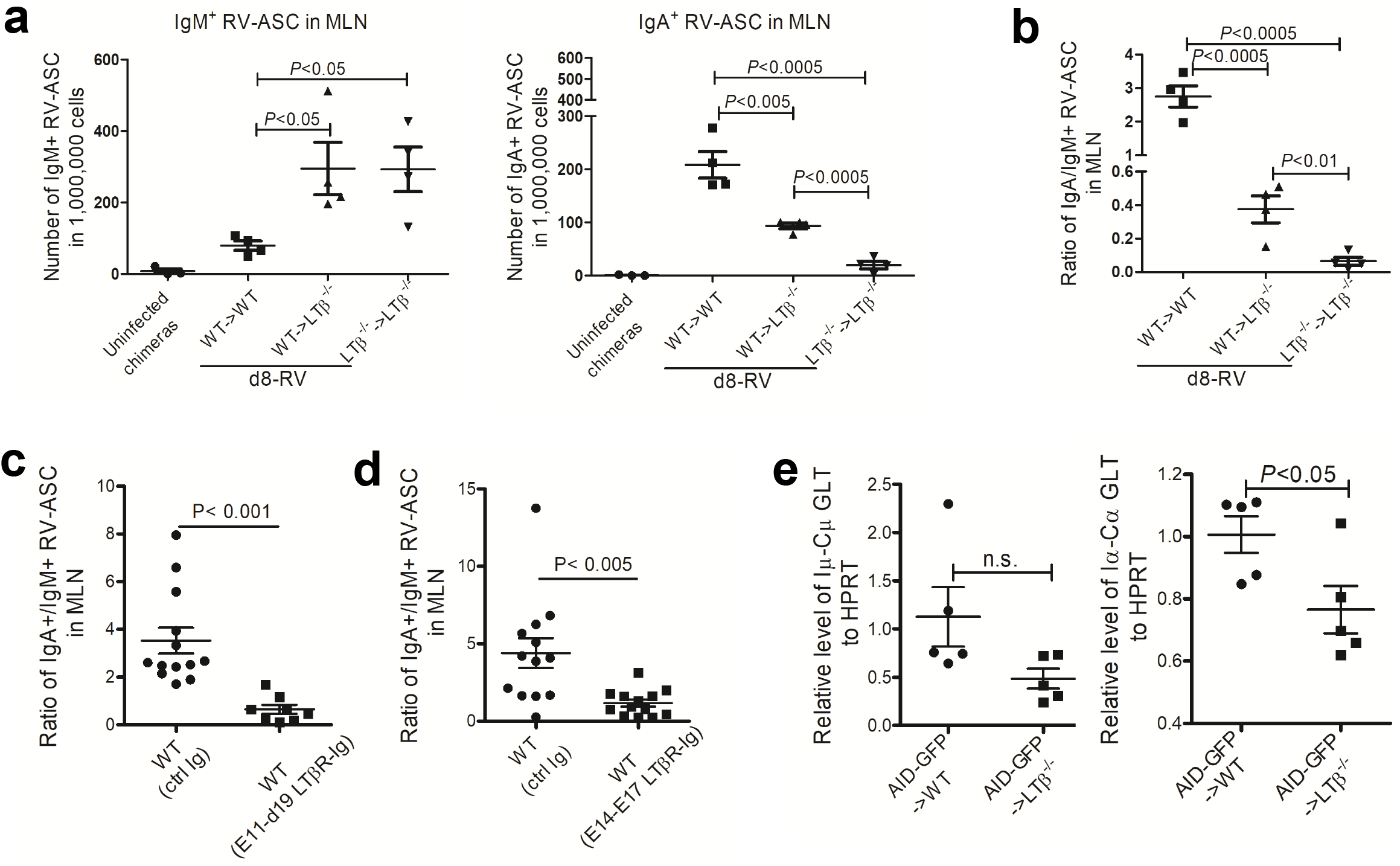
Early life LTβR signalling is required for IgA class switch in the adult MLN. **(a)** Enumeration of IgM+ RV-ASC (left panel) and IgA^+^ RV-ASC (right panel) in the MLN of BM chimeric mice at d8 post-infection. **(b)** The ratio of IgA^+^ *versus* IgM+ RV-ASC in the MLN of chimeric mice at d8 post-infection. Data in (a-b) represent two independent experiments (n=3-4 mice per group). **(c)** The ratio of IgA^+^ *versus* IgM+ RV-ASC in in the MLN of WT mice that received E11-d19 LTβR-Ig *versus* control Ig at d8 post-infection. **(d)** The ratio of IgA^+^ *versus* IgM+ RV-ASC in in the MLN of WT mice that received E14/E17 LTβR-Ig *versus* control Ig at d8 post-infection. Data in **(c-d)** were pooled from 3 independent experiments (n= 8-13 mice per group). **(e)** Levels of Iμ-Cμ and Iα-Cα germline transcripts in AID-GFP+/B220+ cells isolated from the MLN of AID-GFP➔WT or AID-GFP➔LTβ^-/-^ chimeras at d8 post-infection.

To further characterize the phenotype of ASC generated in the MLN of LTβ^-/-^ chimeras, we next assessed the capacity for B cells to upregulate Blimp-1, a transcription factor that is essential for plasma cell differentiation (*36, 37*). To test this, Blimp-1^YFP^ BM was transferred into WT and LTβ^-/-^ hosts, and the expression of Blimp-1 was tracked in MLN-resident B cells at d8 post-RV infection (**Supplementary Fig. 5a**). We observed that the induction of YFP in MLN B cells was normal in Blimp-1^YFP^ ➔LTβ^-/-^ chimeras when compared to Blimp-1^YFP^➔WT controls (**Fig. 4a-b**), However, because the reduction in frequency of IgA^+^ RV-specific ASC was also quite pronounced in the SILP of mice that lack LTβR signalling *in utero* (**Fig. 1c and 1l**), we hypothesized that perhaps subsequent maturation of Blimp-1+RV-induced PC may be impaired in WT➔LTβ^-/-^ chimeric mice. Since the differentiation of IgA^+^ PC involves the progressive downregulation of B220, CD19 and MHC II molecules and upregulation of CCR10 (*19, 37-39),* we used low B220 levels as a distinguishing factor for late-stage PC. RNA-sequencing analysis of B220^high^YFP^high^ and B220^low^YFP^high^ B cell subsets from RV-immunized Blimp-1^YFP^ mice validated that B220^low^YFP^high^ cells have a phenotype consistent with that of late-stage PC. Specifically, elevated expression of *Ccr10* concomitant with reduced levels of *Cd19* and MHCII molecules *H2-Aa, H2-Ab1, H2-DMb2* and *H2-Ob* was observed in B220^low^YFP^high^ B cells compared to B220^high^YFP^high^ B cells, consistent with a late-stage PC phenotype (**Fig. 4c**) (*19, 37-39*). Using this strategy, we assayed expression of genes such as *Ccr9, Ccr10* and *Itgb7,* all of which are involved in IgA^+^ ASC migration to the SILP. We found that the mRNA levels of all of these genes were significantly reduced in the B220^low^YFP^high^ B cells from the MLN of Blimp-1^YEP^➔LTβ^-/-^ chimeras at d8 post-immunization when compared to control WT chimeras (**Fig. 4d**), indicating that the phenotype of B220^low^YFP^high^ B cells is altered in mice where LTβ expression is absent in early life.

**Fig. 4.**
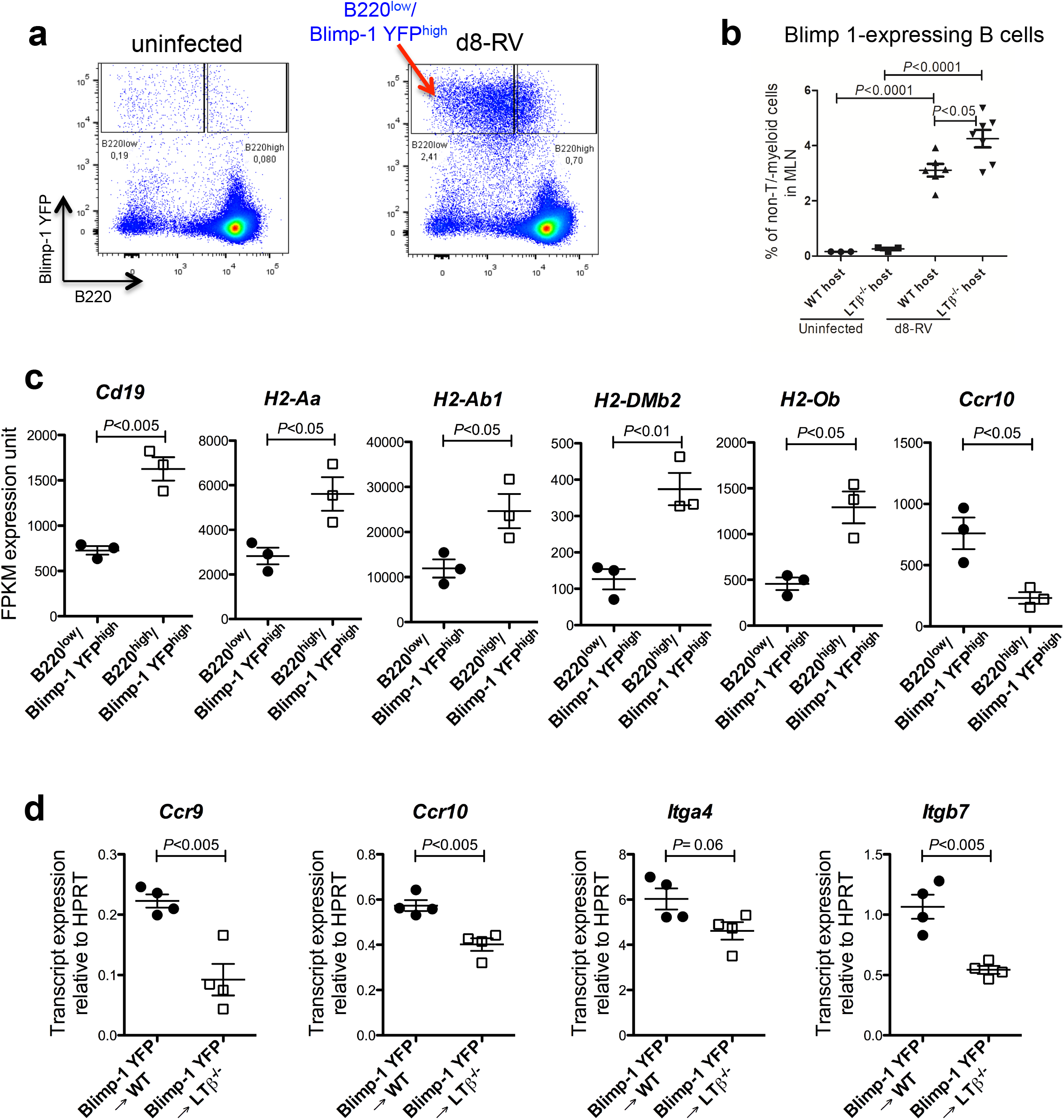
Perinatal LTβR signalling is required for the accumulation of gut-homing plasma cells in adults. **(a)** Representative FACS plots of B220^low^YFP^high^ and B220^high^YFP^high^ B cells derived from the MLN of uninfected *versus* d8 post-infection Blimp-1^YFP^➔WT chimeric mice. **(b)** Frequency of YFP^high^ B cells in the MLN of Blimp-1^YFP^➔WT *versus* Blimp^-/-^➔LTβ^-/-^ chimeric mice. Data represent at least four independent experiments each with three to five mice per group. **(c)** Whole genome RNA-sequencing of B220^low^YFP^high^ and B220^high^YFP^high^ B cells sorted from MLN of Blimp-1^YFP^➔WT chimeric mice at d8 post-infection. n= 3 samples per group, and each sample is pooled from 2-3 mice. **(d)** Digital droplet PCR analysis of indicated genes in B220^low^YFP^high^ B cells sorted from MLN of Blimp-1^YFP^➔WT *versus* Blimp-1^YFP^➔LTβ^-/-^ chimeric mice at d8 post-infection. n= 4 samples per group, and each sample is pooled from 2-3 mice.

In summary, although early life expression of LTβ is not required for the initiation of a RV-driven GC response and AID induction in adult MLN B cells, it is required for CSR to IgA and the upregulation of gut-homing markers on Blimp-expressing B220^low^ B cells.

### LTβR signalling *in utero* dictates the phenotype of MLN-resident stromal cells during adulthood

To test whether the anti-RV IgA CSR defect in WT➔LTβ^-/-^ chimeras is B cell intrinsic or extrinsic, we sorted MLN B cells from uninfected WT➔WT and WT➔LTβ^-/-^ chimeras and induced IgA and IgG1 CSR *ex vivo.* We found that MLN B cells derived from LTβ^-/-^ chimeras exhibited no CSR defect *ex vivo* (**Fig. 5a**). Therefore, consistent with the notion that LTβR is not expressed in B cells but rather in DC, macrophages and lymphoid stromal cells (including follicular DC) (*4, 6, 7*), the defect in CSR we observe in the MLN of mice that lack early life LTβR signalling is B cell-extrinsic.

**Fig. 5.**
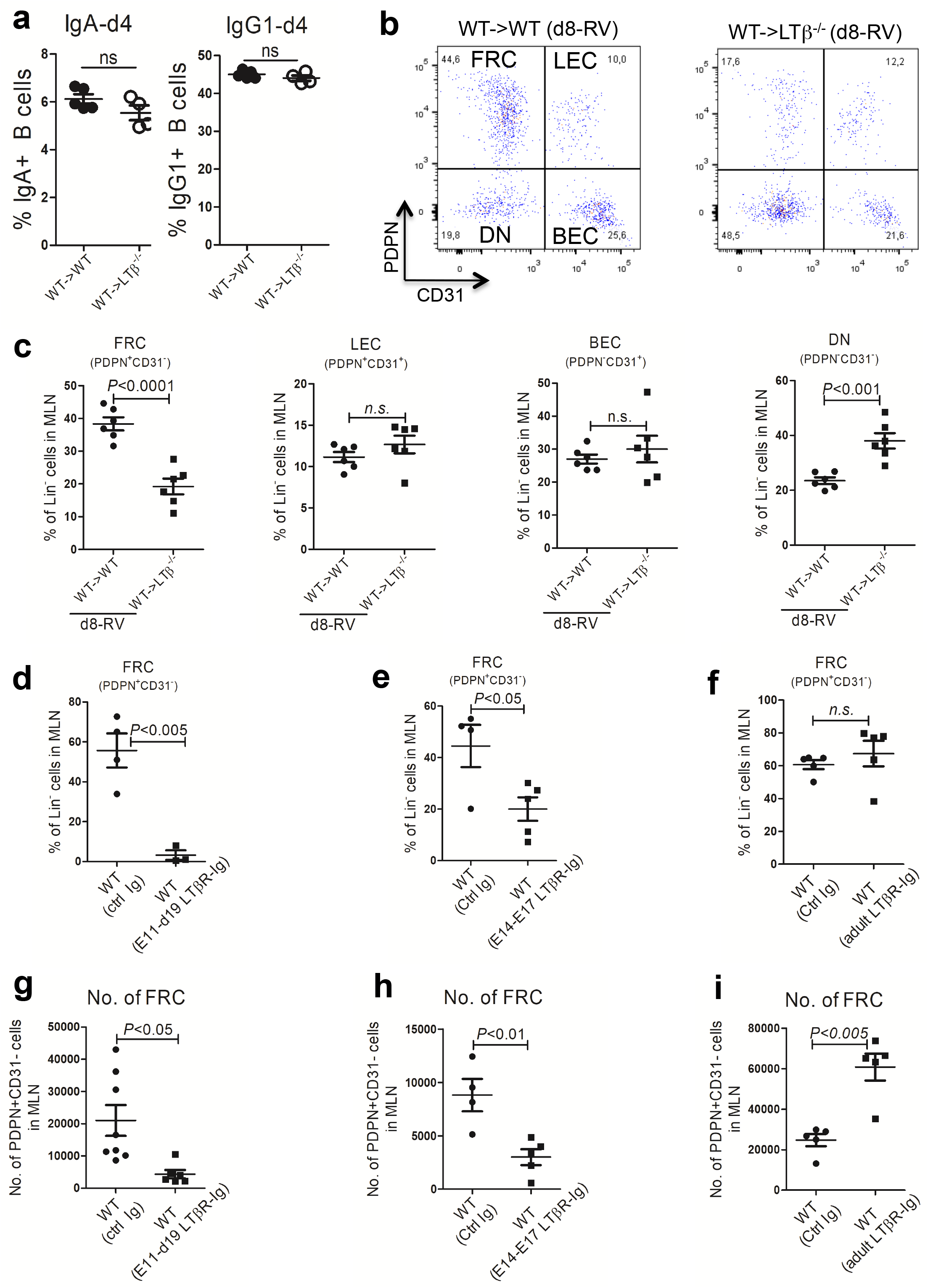
Early life LTβR signalling dictates the phenotype of lymphoid stromal cells in adult MLN. **(a)** Frequency of IgG1 and IgA *ex vivo* class switched B cells derived from the MLN of naïve WT➔WT or WT➔LTβ^-/-^ chimeric mice (n=4-5 mice per group). Data represent two independent experiments. **(b)** Representative flow cytometry plots of MLN stromal cells (FRC: fibroblastic reticular stromal cells; LEC: lymphatic endothelial cells; BEC: blood endothelial cells; DN: PDPN/CD31 double negative stromal cells). **(c)** Frequency of FRC among lineage-negative (Lin^−^) stromal cells in the MLN comparing WT➔WT and WT➔LTβ^-/-^ chimeric mice at d8 post-infection (n= 6 mice per group). Data represent two independent experiments. **(d-f)** Frequency of PDPN+CD31^−^ FRC among Lin^−^ cells in the MLN of WT mice that received LTβR-Ig or control Ig at E11-d19 **(d)**, at E14/17 **(e)**, or during adulthood **(f)** (n=3-5 mice per group). **(g)** Absolute numbers of PDPN+CD31^−^ FRC among Lin^−^ stromal cells in the MLN of WT mice that received LTβR-Ig or control Ig at E11-d19. **(h)** Absolute numbers of PDPN+CD31^−^ FRC among Lin^−^ stromal cells in the MLN of WT mice that received LTβR-Ig or control Ig atE14/17. **(i)** Absolute numbers of PDPN+CD31^−^ FRC among Lin^−^ stromal cells in the MLN of WT mice that received LTβR-Ig or control Ig during adulthood. (n=3-8 mice per group). Data in **(d-i)** represent at least two independent experiments. No. = number.

Since it has been demonstrated that lymphoid stromal cells play an important role in modulating B cell and plasma cell responses in peripheral lymphoid tissues (*37, 40-42*), we reasoned that stromal cell-intrinsic LTβR signalling *in utero* may be important for generating IgA^+^ RV-specific ASC within the adult MLN. To test this, we first examined the expression of podoplanin (PDPN) and CD31 on lineage-negative (lin^−^) MLN stromal cells in WT *versus* LTβ^-/-^ adult mice infected with RV in order to identify PDPN+/CD31^−^ fibroblastic reticular stromal cells (FRC), PDPN+/CD31+ lymphoid endothelial cells (LEC), PDPN^−^CD31^+^ blood endothelial cells (BEC) and PDPN^−^CD31^−^ double negative cells (DN) (*43*) (see **Supplementary Fig. 6a**). Compared to d8-RV infected WT mice, the frequency of FRC among lin^−^ cells was significantly reduced in the MLN of LTβ^-/-^ mice, while the representation of DN cells was proportionally increased (**Supplementary Fig. 6b-c**). We did not calculate absolute numbers of FRC in MLN of LTβ^-/-^ mice, as the decreased cellularity in LTβ^-/-^ MLN would likely skew the analysis (as opposed to mice receiving LTβR-Ig *in utero* – see below).

Next we examined the effect of early life LTβR signaling on FRC in the inflamed MLN post-RV infection in chimeric mice. In comparison to LTβ^-/-^ mice, we observed a similarly disproportionate loss in FRC in the MLN of WT➔LTβ^-/-^ chimeras compared to WT➔WT controls at d8 post-RV infection, indicating that skewing towards DN stromal cells in the MLN could not be rescued with the transplantation of WT BM (**Fig. 5b-c**). This observation was recapitulated in mice treated with LTβR-Ig from E11-d19 (**Fig. 5d and 5g**), and from E14/E17 (**Fig. 5e and 5h**), but not when LTβR-Ig was administered in adulthood (**Fig. 5f and 5i**). Moreover, LTβR-Ig treatment at E14/E17 reduced the expression of MLN stromal cell activation markers (*44*), including VCAM-1 and MHC class I (H2-Kb) (**Supplementary 7a-c**). Next we employed RNA-sequencing on purified MLN stromal cells from mice that were treated at E14/E17 with LTβR-Ig *versus* control Ig. Among the known positive regulators of gut IgA responses (*45-49*), we found a significant reduction in *Mmp2*, *Mmp9*, *Cxcl12*, *Il33*, *Il1b* and *Tnfsf12* transcripts and a trend towards reduced *Tnfsf13b* (*Baff*) and *Il6* transcripts in mice that received LTβR-Ig *in utero* (E14/E17) (Supplementary 7d-e). In summary, LTβR signalling *in utero* influences the phenotype and gene expression profile of stromal cells in MLN of adult mice

### Deletion of LTβR signalling in stromal cells impairs mucosal anti-RV IgA responses

Because a reduction in FRC was observed in the adult MLN of mice treated *in utero* with LTβR-Ig, and because these stromal cells exhibited defects in many pro-IgA switch factors, we hypothesized that specific deletion of LTβR in FRC would result in an impaired IgA response to RV in the adult mouse. To test this, we examined the RV-specific IgA response in *Ltbr^fl/fl^* x Ccl19-cre mice which lack LTβR signalling specifically in FRC (*50*). Consistent with previous studies examining the inguinal LN (*50*), the frequency of FRC in the MLN of *Ltbr^fl/fl^* x Ccl19-cre mice among lin^−^ stromal cells was significantly reduced when compared to littermate controls (**Fig. 6a-b**). Importantly, unlike mice treated with LTβR-Ig at E14/17 or WT➔LTβ^-/-^ chimeric mice, *Ltb^fl/fl^* x Ccl19-cre retain the majority of their PP (**Fig. 6c**) thus allowing us to definitely disentangle a putatitive role for PP in the anti-RV IgA immune response. Importantly, when we examined the RV response in *Ltb^fl/fl^* x Ccl19-cre mice we observed a significant reduction in fecal anti-RV IgA titres when compared to littermate controls (**Fig. 6d**). Moreover, compared to littermate controls we observed a defect in IgA CSR in the MLN and a corresponding reduction in the accumulation of IgA^+^ RV-ASC in the SILP in *Ltbr^fl/fl^* x *Ccl19*-cre mice (**Fig. 6e-f**), further demonstrating that LTβR signalling in stromal cells is required for the induction and/or maintenance of an intestinal anti-RV IgA response. In summary, these data provide direct evidence that LTβR expression in FRC is required for the anti-RV IgA response.

**Fig. 6.**
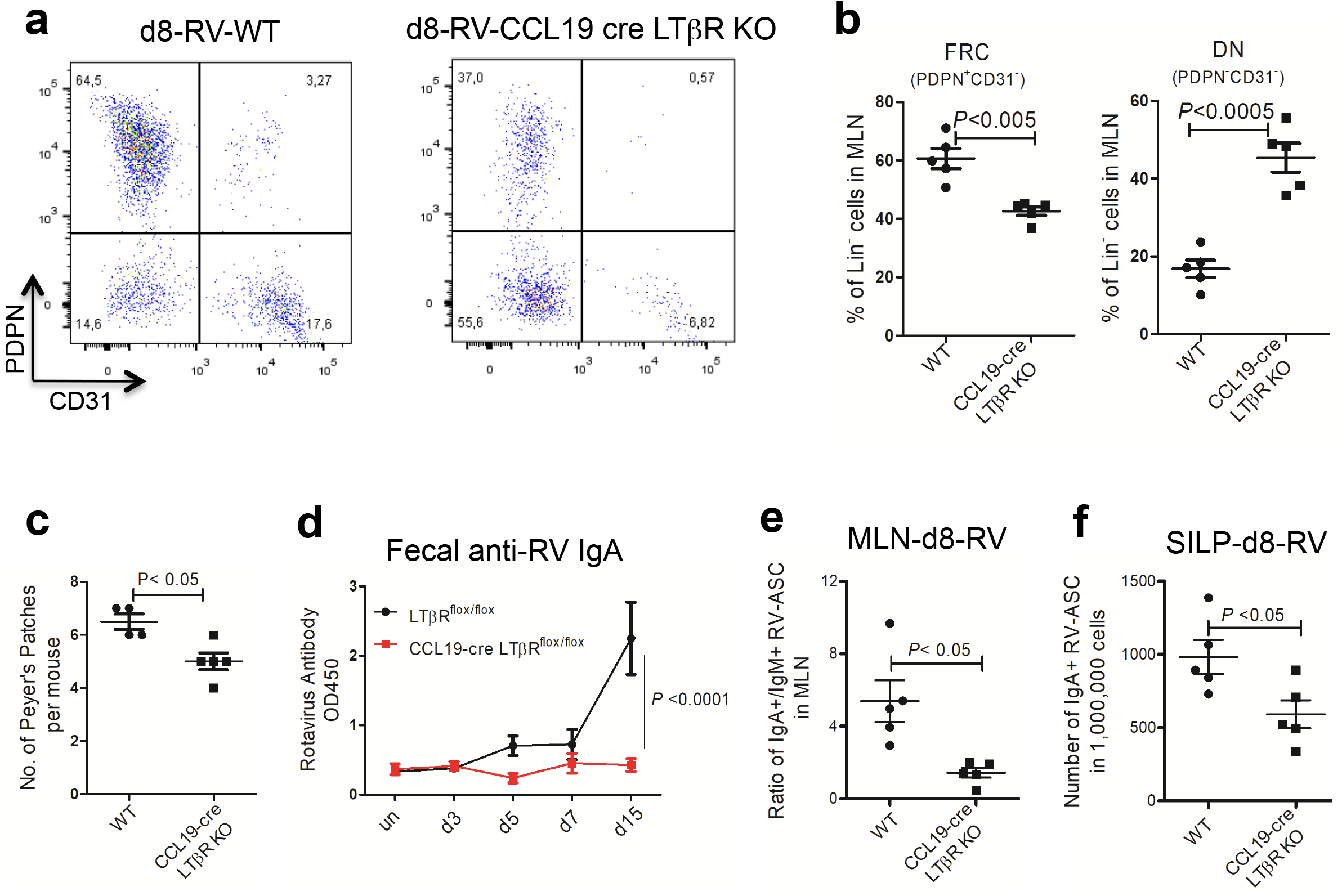
Deletion of LTβR in FRC impairs the anti-RV IgA response. **(a)** Representative FACS plots of MLN stromal cells from CCL19-cre/LTβR^flox/flox^ *versus* WT littermate controls at d8 post-infection. **(b)** Frequency of FRC and DN stromal cells among Lin^−^ cells was compared between d8-RV infected CCL19-cre/LTβR^flox/flox^ and WT littermate controls. Data are representative of three independent experiments (n=3-5 mice per group). **(c)** Number of Peyer’s Patches per mouse in d8-RV infected CCL19-cre/LTβR^flox/flox^ and WT littermate controls (n=3-5 mice per group). **(d)** Fecal anti-RV IgA levels in CCL19-cre/LTβR^flox/flox^ *versus* WT littermate control mice followed at the indicated time points post-infection (n= 4-5 mice per group). Data were analyzed by two-way ANOVA. **(e)** Ratio of IgA^+^ *versus* IgM+ RV-ASC MLN of CCL19-cre/LTβR^flox/flox^ *versus* WT littermate controls at d8 post-infection. **(f)** Enumeration of IgA^+^ RV-ASC in the SILP of CCL19-cre/LTβR^flox/flox^ *versus* WT littermate controls at d8 post-infection. Data in **(e-f)** represent three independent experiments (n=3-5 mice per group).

## DISCUSSION

In this study we discovered a novel role for LTαβ/LTβR signalling during the *in utero* period in shaping the humoral immune response to RV infection during adulthood. We further show that early life LTβR signalling shapes MLN stromal cell gene expression and phenotype. Consequently, while the initiation of a GC response and upregulation of AID within B cells in the adult MLN is normal, subsequent class switch to IgA and the acquisition of gut homing molecules on post-GC Blimp1+ PC is strongly affected. Lastly, we found that deletion of *Ltbr* in FRC recapitulates this phenotype. We postulate that the combination of a reduction of FRC as well as diminished expression of key pro-IgA factors in remaining FRC in the MLN of mice that lack LTβR *in utero* accounts for the pronounced IgA CSR defect in response to RV infection in these mice.

PP have been thought as the major site for homeostatic commensal-specific IgA induction and for initiating the intestinal immune response to pathogens (*51*). Antigen and immune cells from PP can reach the MLN via lymphatics, with MLN serving as a second line of defense (*52, 53*). Mice that lack LTβ *in utero*, such as those we studied herein, lack PP (*10*). This could potentially complicate the interpretations of our results. Nevertheless, we found that during RV infection, the induction of AID-GFP expression, and the generation of IgA^+^ RV-specific ASC were focused within the MLN rather than the PP. This is consistent with work from Greenberg and colleagues who observed that IgA^+^ RV-ASC peaked in the MLN at approximately day 7 postinfection compared to the PP at day 270 post-RV infection, suggesting that during the early phases of the RV IgA response, generation of RV-specific IgA ASC occurs largely in the MLN rather than the PP (*20*). It could also be argued that the PP helps to shuttle RV antigen into the MLN where the immune response is then initiated, and indeed, RV has been shown to localize to the dome of the PP (*29*). However, in the MLN of *in utero* LTβR-Ig treated mice there is clearly enough RV antigen to induce an anti-RV IgM response and also sufficient antigen to induce AID and Blimp-1 expression in MLN-resident B cells even though these mice lack PP, suggesting that RV may access gut-draining MLN via alternative mechanisms in the absence of PP. Moreover, in mice where LTβR is specifically deleted in FRC (*Ccl19*-cre x *Ltbr*^fl/fl^), PP are present yet the RV-specific IgA response is markedly impaired. Thus, PP are not required for induction of an anti-RV IgA response, at least in the first weeks following RV infection. Taken together, the defects in the anti-RV IgA response that we observe in mice that lack LTβ signalling *in utero* are likely not a result of the absence of PP.

It has been previously shown that DC-intrinsic LTβR signalling is required for polyclonal anti-commensal IgA responses generated in the PP (*21*). Our previous studies found that DC- and macrophage-intrinsic LTβR signalling are dispensable for anti-RV IgA responses (*54*). PP harbour distinctive types of DC that may play unique roles in inducing anti-commensal IgA responses in this location (*55, 56*). Since we show that the anti-RV response is largely focused to the MLN, this could explain why the anti-RV IgA response does not require myeloid cell-intrinsic LTβR expression.

In contrast to anti-RV IgA responses, we found that polyclonal IgA responses do not require early life LTαβ/LTβR signalling. Anti-commensal polyclonal B cell responses are thought to be chronic, and largely independent of T cell help (*57*). In contrast, we and others have found that the anti-RV IgA response is largely T-dependent (**Supplementary Fig. 7f-g**) (*58*). We hypothesize that this differential reliance on T cell help could account for why anti-RV IgA responses, but not polyclonal (anti-commensal) IgA responses, are dependent on early life LTβR signalling.

MLN are preserved in LTβ^-/-^ mice and in mice treated from E14 with LTβR-Ig, thus allowing us to study the role of perinatal LTαβ/LTβR signalling on the initiation of the RV response. One potential confounder is that, even though MLN develop in the absence of LTβ, they may be entirely incapable of initiating a GC response. This notion is countered by studies performed by Flavell and colleagues several years ago where they noted the presence of GC in the MLN of LT-deficient mice (*59*). In agreement, we also found that the initiation of the humoral immune response to RV in terms of the formation of GC B cells, the upregulation of AID (as reported by AID-GFP expression) and the upregulation of Blimp-1 (as reported by YFP expression), were all normal in the MLN of LTβ^-/-^ mice transplanted with WT BM and/or WT mice treated *in utero* with LTβR-Ig. This suggests that the structure of the LTβ^-/-^ MLN can accommodate the early steps of the anti-RV B cell response. However subsequent steps of this response, namely the expression of Iα-Cα germline transcripts, class switch to IgA, accumulation of Blimp1+B220^low^ B cells and the expression of *Ccr9, Ccr10* and *Itgb7* in Blimp1+B220^low^ B cells, are all impaired in the MLN of LTβ^-/-^ mice reconstituted with WT BM and/or mice treated *in utero* (E14/E17) with LTβR-Ig. These defects are accompanied by a dramatic reduction in RV-specific IgA^+^ ASC in the SILP and RV-specific IgA in the feces. Therefore, while the GC response occurs in the LTβ^-/-^ MLN, the capacity of the GC to support class switch, and the quality of the emerging Blimp-1+ cells, including their ability to reach the SILP effector site, is significantly deranged.

TGFβ is a very key pro-IgA factor (*60*), thus the impaired induction of Iμ and Iα germline transcripts in the absence of early life LTβR signalling might be due to limited availability of the active cleaved form of TGF-β1 within the MLN microenvironment. Indeed, *Mmp2* and *Mmp9,* which encode the matrix metalloproteinases that cleave TGF-β1 from its latent complex (*46*) were both markedly reduced in MLN stroma after *in utero* (E14/E17) inhibition of LTβR signalling. In addition to these MMPs, the expression of well-known pro-IgA switch factors *Tnfsf13b* and *Il6 (47*), as well as *Il33*, *Il1b* and *Tnfsf12*, were all reduced in MLN stromal cells derived from mice that received LTβR-Ig *in utero* (E14/E17). IL-33 has been shown to synergize with TGF-β to promote IgA generation and mice deficient in IL-33 have much lower levels of intestinal IgA (*45*). IL-1β-deficient mice also manifest decreased levels of IgA-producing cells in the SILP and reduced levels of intestinal IgA, concomitant with decreased mRNA expression of inducible nitric oxide synthase (iNOS) in the small intestine (*48*). It has been reported that patients with autosomal dominant deficiency in *Tnfsf12* exhibit reduced levels of IgA, although the molecular mechanisms by which TNFSF12 regulates IgA production are unclear (*49*). Collectively, we reason that a putative reduction in active TGF-β1 along with a reduction in other pro-IgA switch factors expressed by MLN stromal cells, conspires to create an environment in the adult MLN that fails to promote IgA class switch and the proper upregulation of mRNA encoding LP homing molecules *(Ccr9, Ccr10, Itga4, Itgb7) (19, 20*). This “compound” FRC defect results in a profound reduction in RV-specific IgA production in the gut, and to our knowledge, this is the first example of a molecular signal during fetal life that imprints the phenotype of MLN-resident FRC in the adult. In contrast, although some stromal cells such as follicular DC are highly sensitive to LTβR-Ig treatment in the adult mouse (*61*), treatment of adult mice with LTβR-Ig did not affect the frequency of FRC proportional to other lin^−^ MLN cells, nor the IgA response to RV. It remains to be investigated whether the *in utero* LTβR signalling-dependent stromal cells we have implicated in the IgA response to RV are involved in the immune response against other intestinal pathogens.

Early life exposures including the maternal microbiota and microbiota-associated metabolites, maternal nutrition and maternal antibodies have all been shown to have important effects on the development of the mucosal and peripheral immune system (*62*). These exposures may ultimately influence immune responses to vaccines. Indeed, vaccination against RV, the leading cause of childhood diarrhea worldwide (*63*), works well in North American children, yet is less efficacious in resource-poor countries (*64, 65*). Identifying molecular pathways that operate in the perinatal period to poise the organism for an optimal immune response to pathogens later in life (such as the LT pathway) will help inform vaccine design. Moreover, in addition to IgA production, several recent studies including our own have shown that IgA-producing PC have other functions, notably the production of cytokines and the expression of checkpoint inhibitory molecules that play immunosuppressive roles in the immune system (*66-69*). Thus, an understanding of how the MLN environment fosters the generation and function of IgA-producing B cells is important for gaining insights into mechanisms of immune system homeostasis.

## MATERIALS AND METHODS

### Mice

C57BL/6 WT mice (Charles River Laboratory, St. Constant, QC, Canada) were bred in the Division of Comparative Medicine Animal Vivarium facility at the University of Toronto. LTβ^-/-^ mice were originally from B&K Universal and bred in our animal facility (*70*). AID-GFP mice were obtained from Dr. Rafael Casellas, and Blimp-1^YFP^ mice were purchased from the Jackson Laboratory. Bone marrow chimeras were generated as we previously described (*34, 71*); irradiated mice were rested for 8-12 weeks to allow for bone marrow reconstitution, and were kept on neomycin sulfate water (2g/L) for the first two weeks. The mice were maintained under specific pathogen-free conditions. The experimental procedures were approved by the Animal Care Committee of University of Toronto and by the Cantonal Veterinary Office (St. Gallen, Switzerland).

### Tissue harvest and cell isolation

At the indicated time points, all MLN or PP were harvested and ground between glass slides, followed by filtration with a 70-μm cell strainer. BM cells were flushed out from femurs and tibia of mice followed by red blood cell lysis. Single cell suspensions from PP were prepared by grinding PP between glass slides, followed by filtration with a 70-μm cell strainer. For AID-GFP FACS and RV-ELISPOT with SILP cells, PP were first removed from the small intestine. The segmented small intestine was washed with CMF buffer (HBSS +2% FBS+ 15mM HEPES), and then vigorously shaken in CMF/EDTA buffer (5mM EDTA), followed by digestion with freshly made enzyme mix comprised of RPMI-1640 containing Collagenase IV (0.25mg/mL; Sigma) and DNase I (0.025mg/mL; Roche) in a 37°C water bath.

### Fecal polyclonal IgA ELISA

Fecal pellets were collected from the indicated groups at approximately 2 months following BM transplantation. Fecal supernatant was prepared (10% wt/vol) with PBS. The Ig isotype-specific ELISA assays were performed as we previously described (*72*). Briefly, 96-well ELISA plates (NUNC) were coated overnight at 4°C with goat anti-mouse Ig (SouthernBiotech). After blocking with 2% BSA, 1/5 diluted fecal samples were added into the wells, followed by serial 3-fold dilutions. The bound antibodies from fecal supernatants were detected with goat antimouse IgA that was conjugated with alkaline phosphatase (SouthernBiotech), followed by development with TMB substrate (Bioshop).

### RV-specific ELISA and ELISPOT

WT mice, AID-GFP mice or BM chimeric mice were infected with RV as we previously described (*54, 73*). Fecal pellets were collected from each mouse one day before RV challenge and on the indicated days post-infection. Fecal supernatant was prepared (10% wt/vol) with PBS, and ½ dilution of fecal supernatant was used for anti-RV IgA and RV antigen ELISAs, as we previously described (*54, 73*). For anti-RV IgA ELISA, serial dilutions of RV infected samples were used to validate the assay, and all the samples shown in the individual graph were tested in one ELISA plate. At the indicated days post-RV infection, an RV-specific ELISPOT assay was performed, measuring the numbers of RV-specific IgA^+^ or IgM+ ASC from indicated tissues. Lymphocytes from the SILP were prepared by Percoll gradient as previously described (*54*). In brief, MultiScreen-HTS-HA filter plates were coated overnight with inactive antigen (Microbix), and then blocked with complete RPMI media. The plates were incubated with two-fold serial dilutions of MLN, BM or SILP lymphocytes overnight, followed by detection with HRP-conjugated goat anti-mouse IgA (SouthernBiotech). AEC (Vector Laboratories) was then used as the colorimetric precipitating substrate for HRP to develop the plate. Positive spots on the membrane within each well were counted in a blinded manner with a Nikon stereomicroscope.

### Pharmacological inhibition of LTβR signalling

To pharmacologically inhibit LTβR signalling in adult mice, WT mice were intraperitoneally injected with 100μg LTβR-Ig or isotype control antibodies on the indicated days. To inhibit LTβR signalling *in utero,* pregnant WT dams were injected with 200μg LTβR-Ig or control antibodies at day 14.5 and 17.5 of pregnancy via tail vein injections, as previously described (*35*). The E14/17 LTβR-Ig treated pups were co-caged with control Ig treated pups since weaning (i.e. 3 weeks old). To inhibit LTβR signalling during both fetal life and neonatal periods, pregnant WT dams were injected with 200μg LTβR-Ig or control antibodies at d11.5, 14.5 and 17.5 of pregnancy via tail vein injections, and the delivered offspring received injections of LTβR-Ig or control antibodies (100μg antibody per 20 gram of mice) at neonatal days 5.5, 12.5 and 19.5. The E11-d19 LTβR-Ig treated pups were co-caged with control Ig treated pups since weaning.

### MLN Blimp-1+ B cell flow cytometry and flow cytometric sorting

Single cell suspensions from the draining MLN of Blimp-1^YFP^ mice or chimeric mice were prepared at the indicated time points. Cells were stained with antibodies against mouse CD45.2 (APC-eFluor780, clone: 104), B220 (PE or BV605, clone: RA3-6B2) and APC-dump antibodies (CD3, clone: 17A2; CD11c, clone: N418; F4/80, clone: BM8) in the presence of Fc block (2.4G2). The stained cells were then incubated with 7-AAD (live/dead marker) before FACS acquisition on an LSR II, LSR X-20 or Influx sorter (BD Biosciences). B cells of interest were identified as live CD45^+^/CD3^−^CD11c^−^/F4/80^−^B220^low^/YFP^high^ (late-stage plasma cells) or live CD45^+^/CD3^−^./CD11c^−^/F4/80^−^B220^high^/YFP^high^ (early-stage plasma cells). FACS data were analyzed using FlowJo software (Treestar Inc.). All the above antibodies for FACS staining were purchased from eBiosciences except B220-BV605 which was purchased was from BioLegend.

### Flow cytometry of MLN GC B cells

Flow cytometric analysis of GC B cells was performed as previously described (*21*). In brief, single cell suspensions from MLN of Blimp-1^YFP^ mice or chimeric mice were stained with antibodies against mouse CD45.2 (PE-eFluor610, clone: 104, eBiosciences), PE-Cy7 dump antibodies (CD4, clone: RM4-5; CD8, clone: 53-6.7; CD11c, clone: N418; F4/80, clone: BM8, all from eBiosciences), CD19 (BV605, clone: 6D5, BioLegend), IgD (eF450, clone: 11-26C, eBiosciences), CCR6 (PE, clone: 29-2L17, BioLegend), GL-7 (eF660, clone: GL-7, eBiosciences) and Fas (biotin, clone: Jo2, BD Pharmigen) in the presence of Fc block, and then stained with streptavidin-APC-eFluor 780. The cells were then stained with 7-AAD prior to flow cytometry on an LSR X-20.

### Flow cytometry of MLN Tfh

Prior to staining, single cell suspensions from the MLN were labeled with Live/Dead Aqua (Life Technologies). Cells were then stained with antibodies against mouse CD4 (PE-Cy7, clone: RM4-5), B220 (BV605, clone: RA3-6B2), CXCR5 (biotin, clone: SPRCL5) and PD-1 (APC, clone: J43) in the presence of Fc block, and then stained with streptavidin-APC-eFluor 780. Cells were then permeabilized and fixed using eBioscience perm/fix buffer, followed by intracellular (i.c.) staining with antibody against Bcl-6 (PE, clone: K112-91). Tfh cells were identified as live CD4^+^/B220^−^/CXCR5^+^/PD-1^+^/i.c. Bcl-6^+^. All the above antibodies for FACS staining were purchased from eBiosciences except B220-BV605 which was from BioLegend and Bcl6 which was from BD Pharmigen.

### MLN stromal cell flow cytometry and sorting

MLN stromal cells were isolated using a modified method as previously described (*74*). MLN from indicated groups of mice were dissected into small pieces and digested with freshly made enzyme mix comprised of RPMI-1640 containing 0.12 mg/ml Collagenase P (Roche), 0.4 mg/ml Dispase (Roche) and 0.025 mg/ml DNase I (Roche) in the 37°C water bath for 4 rounds (total digestion time: 45min). After digestion, cells were filtered through a 70μm cell strainer and washed with PBS containing 2% FBS and 3mM EDTA. MLN cells were stained with APC-eFluor 780 dump antibodies (CD45.2, clone: 104; Ter-119, clone: TER-119; EpCAM, clone: G8.8), FITC dump antibody (B220, clone: RA3-6B2), and antibodies against PDPN (PE, clone: 8.1.1) and CD31 (eF450, clone: 390) in the presence of Fc block. The cells were then stained with 7-AAD before FACS acquisition on an LSR X-20 or Influx sorter. All the above antibodies for FACS staining were purchased from eBiosciences.

To assay the activation status of MLN stromal cells, single-cell suspensions of MLN were labeled with Live/Dead Aqua. Cells were then stained with APC-eFluor 780 dump antibodies (CD45.2/Ter-119/EpCAM, eBiosciences), CD19 (FITC, clone: eBio1D3, eBiosciences), PDPN (PE-Cy7, clone: 8.1.1, BioLegend), CD31 (PerCP Cy5.5, clone: MEC13.3, BioLegend), VCAM1 (biotin, clone: 429, BioLegend), H2Kb (MHC-I, APC, clone: AF6-88.5.5.3, BioLegend), and ICAM-1 (BV421, clone: 3E2, BD Biosciences) in the presence of Fc block, and then stained with streptavidin-BV711. The cells were then subject for FACS acquisition on an LSR X-20.

### RNA sequencing

For MLN B cell subset RNA sequencing, each sample was pooled from the 2-3 mice, and 3 samples from 3 independent experiments were submitted for RNA sequencing; for MLN stromal cell RNA sequencing, each sample was pooled from 2-4 mice, and 4 pooled samples from 4 independent experiments were submitted for RNA-sequencing. RNA from the sorted cells was subjected to Illumina Ultra-low Input cDNA library preparation kit, and the subsequent Hi-Seq sequencing was performed in Donnelly Sequencing Center (University of Toronto, Toronto, Canada). RNA sequencing output was subsequently mapped to the mouse whole genome (assembly mm10) using TopHat2 version 2.1.0 in its default setting. The reads mapping to multiple genomic regions were removed from subsequent analysis. The mapped reads were quantified against the refseq annotation using HTSeq-count version 0.7.2 in its default setting. The read counts were normalized using a negative binomial model using Bioconductor package DESeq2 version 1.14.1. The differential expression analysis was performed using DESeq2. Plots using RNA-seq data were generated in R version 3.3.2.

### CD4^+^ T cell depletion

The anti-CD4 InVivoMab (clone: GK1.5, BioXCell) or control antibody was injected intra-peritoneally at 0.5mg/mouse two days prior to RV infection and at d8 post-infection. Blood was collected at day 10 after the second anti-CD4 injection, and stained with FITC-conjugated antimouse CD4 (clone: RM4-5) to verify the depleting efficiency. Mice were sacrificed atd19 post-RV infection, and subjected to RV-specific IgA^+^ ASC analysis.

### Digital droplet PCR

RNA was extracted from CD45^+^/CD3^−^CD11c^−^/F4/80^−^/B220^low^/YFP^high^ cells sorted from MLN using the Qiagen Micro RNA Extraction Kit, followed by cDNA synthesis using Reverse Transcriptase (Thermo Scientific). Expression of genes of interest was measured using 1μL of cDNA with QX200 digital droplet PCR (Bio-Rad). Briefly, using the QX200 automated droplet generator (Bio-Rad), droplets were produced from the mixture consisting of 1μL of cDNA, 227nm of each primer (forward and reverse), 11 μL of Evagreen Supermix (Bio-Rad) and 9μL of ddH_2_O. Once droplets were produced, the samples were subjected to PCR (95 degrees for 5 minutes; 95 degrees for 30 seconds, 61 degrees for 1 minute repeated for 40 cycles; 4 degrees for 5 minutes; 95 degrees for 5 minutes; and infinitely held at 4 degrees). All steps were kept with a ramp rate of 2°C/second. Following PCR, samples were read in the QX200 droplet reader (Bio-Rad). Analysis of droplets was completed using Quantasoft (Version number: 1.7.4.0197) and results were tabulated using GraphPad Prism.

### *Ex vivo* CSR

B cells were sorted from the MLN of co-caged WT➔WT *and* WT➔LTβ^-/-^ chimeras, and IgG1 or IgA isotype switching was induced *ex vivo,* as we previously described (*73*). In brief, single cell suspensions of MLN were stained with anti-B220 (BV605) antibody in the presence of Fc block, and B cell sorting was performed with FACS Aria IIu sorter (BD Biosciences). LN B cells were cultured in complete RPMI with LPS for 4 days. IL-4 was added to induce IgG1 switching, while IL-4, TGF-β, IL-5 and anti-IgD dextran were added for IgA switching.

### Statistical analysis

Data are presented as mean ± SEM. All the data were analyzed using two-tailed Student unpaired t test except for the anti-RV IgA ELISA kinetics that were analyzed by two-way ANOVA, and the frequency of MLN in LTβR-Ig treated mice that were analyzed by chi-square. P<0.05 was considered significant.

## Acknowledgements

We would like to thank Biogen Idec for providing LTβR-Ig and isotype control antibodies, and Dr. Harry Greenberg for providing the murine rotavirus (EC strain). We thank the Gommerman lab members for their advice and discussion during this study, and Dionne White for the help in flow cytometry.

## Funding

This research is supported by a Foundation grant from the Canadian Institutes of Health Research to J.L.G. and a project grant from the Canadian Institutes of Health Research to A.M. Conglei Li is a recipient of the Canadian Institutes of Health Research Postdoctoral Fellowship.

## Author Contributions

C.L. designed the experiments, performed the research, analyzed data, and wrote the manuscript; E.L., C.P., D.L., L.W., A.N., K.K. and T.S. performed the research; M.A. and H.H. analyzed the RNA-seq data; O.R edited the manuscript; A.M. and B.L. analyzed data and edited the manuscript; J.L.G. is the principal investigator who designed the experiments, analyzed data, and wrote the manuscript.

## Competing interests

The authors have no relevant competing interests to declare.

## Data availability

All relevant data are available from the authors.

## Supplementary Figure Legends

**Supplementary Figure 1.**
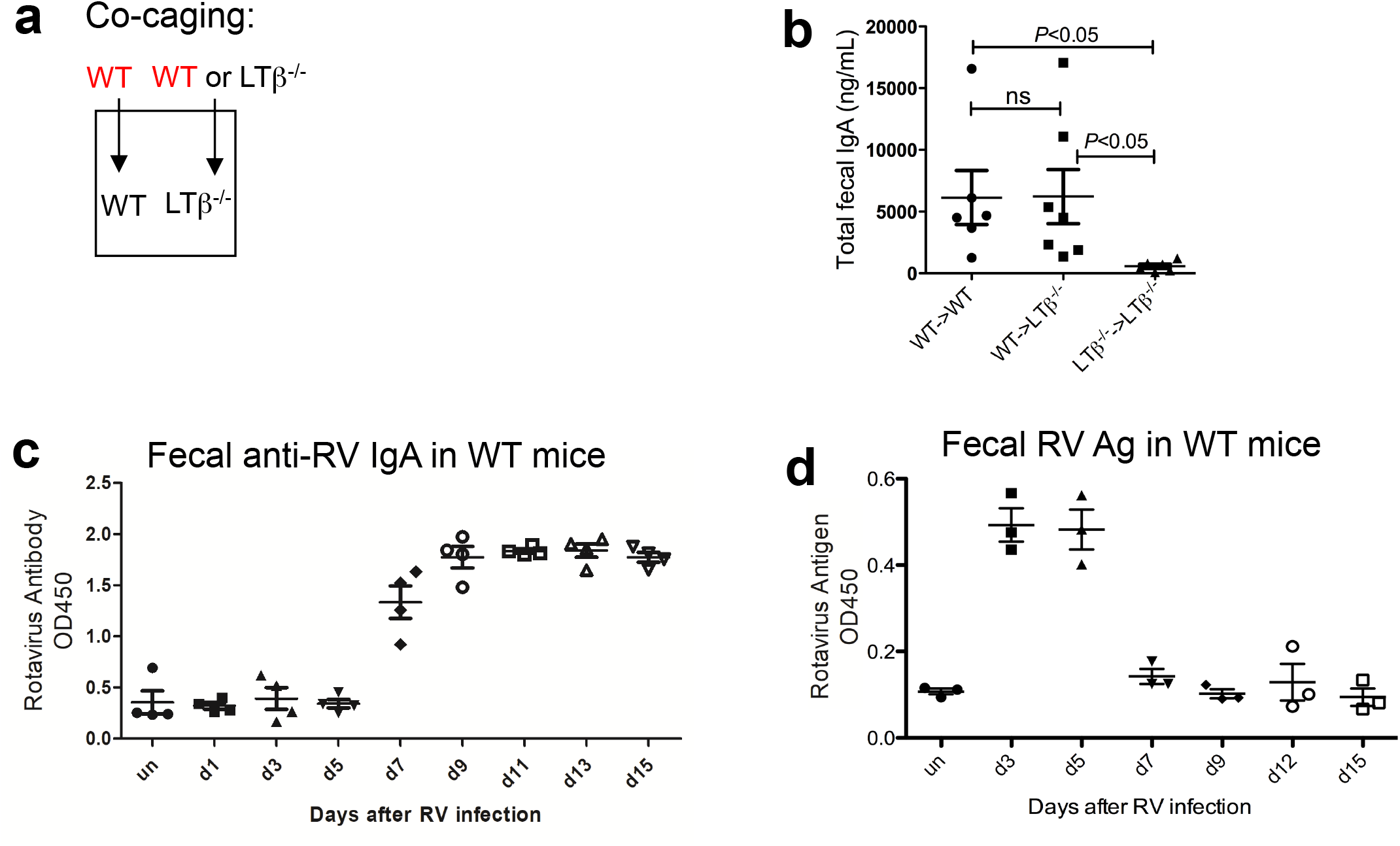
Intestinal polyclonal IgA responses in BM chimeric mice, and kinetics of RV infection in WT mice. **(a)** Diagram depicting co-caging experimental setup for BM chimeric mice. **(b)** ELISA analysis of homeostatic fecal IgA from BM chimeric mice. Data represent two independent experiments each with four to six mice per group. **(c-d)** Measurement of RV-specific IgA and RV antigen in WT mice at various time-points post-infection (n=3-4 mice per group). All samples in **(c)** or **(d)** were tested in one ELISA plate.

**Supplementary Figure 2.**
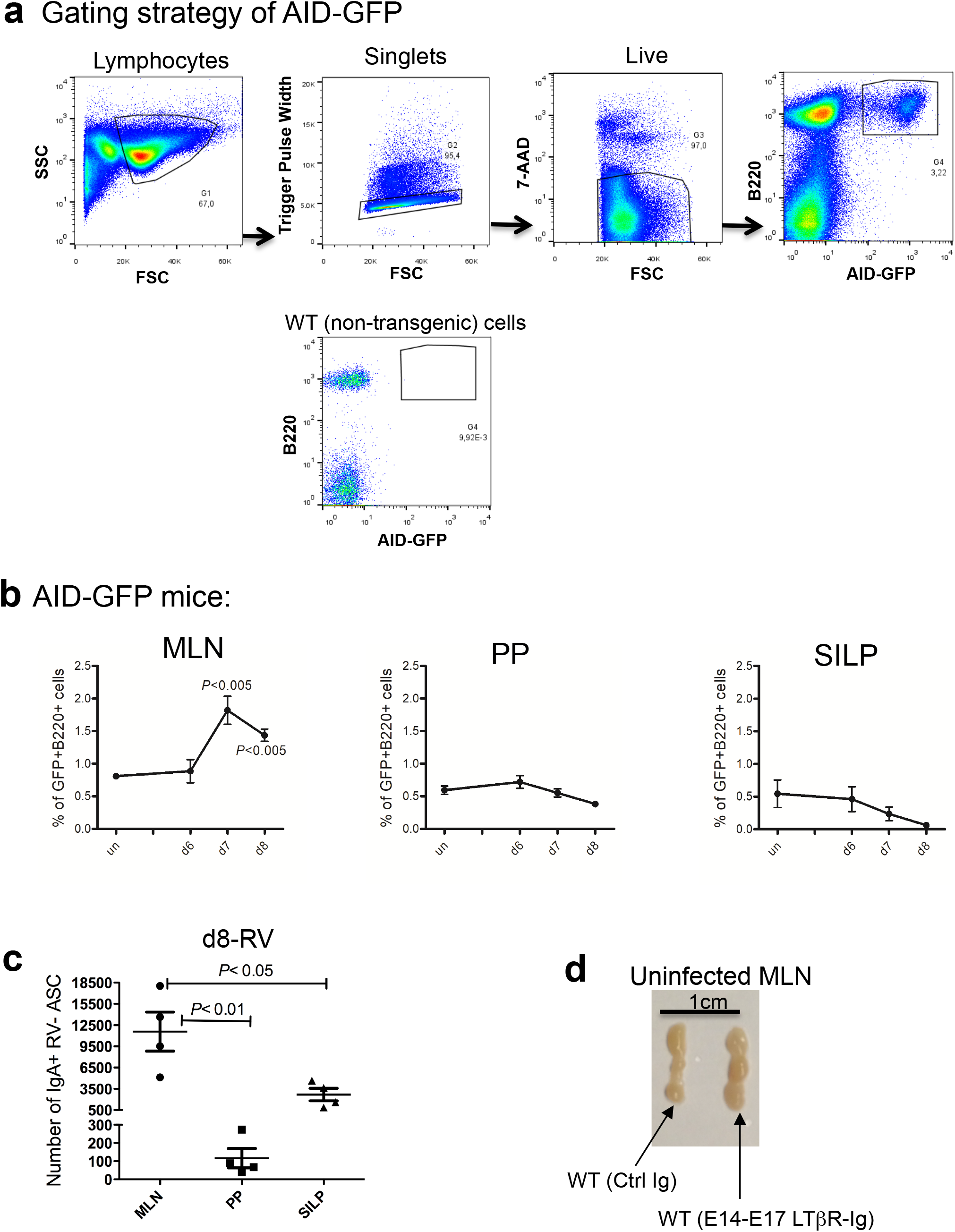
The MLN is the major initiation site of B cell activation in response to RV infection. **(a)** Gating strategy for identification of AID-GFP expressing B cells. Single cell suspensions from the MLN are shown as an example but similar results were obtained in other tissues (i.e. SILP and PP). Background GFP levels in WT mice is shown for gating purpose. **(b)** Graphical representation of the frequency of AID-GFP+/B220+ cells in the indicated tissues during the first 8 days post-infection of AID-GFP mice (un: uninfected). Data represent three independent experiments each with three to four mice per time point. **(c)** Analysis of the number of IgA^+^ RV-ASC in the indicated tissues at d8 post-infection. Data represent two independent experiments each with four mice per group. **(d)** Representative picture of MLN from naïve *in utero* (E14/E17) LTβR-Ig or control Ig treated WT mice (Ctrl Ig: control Ig).

**Supplementary Figure 3.**
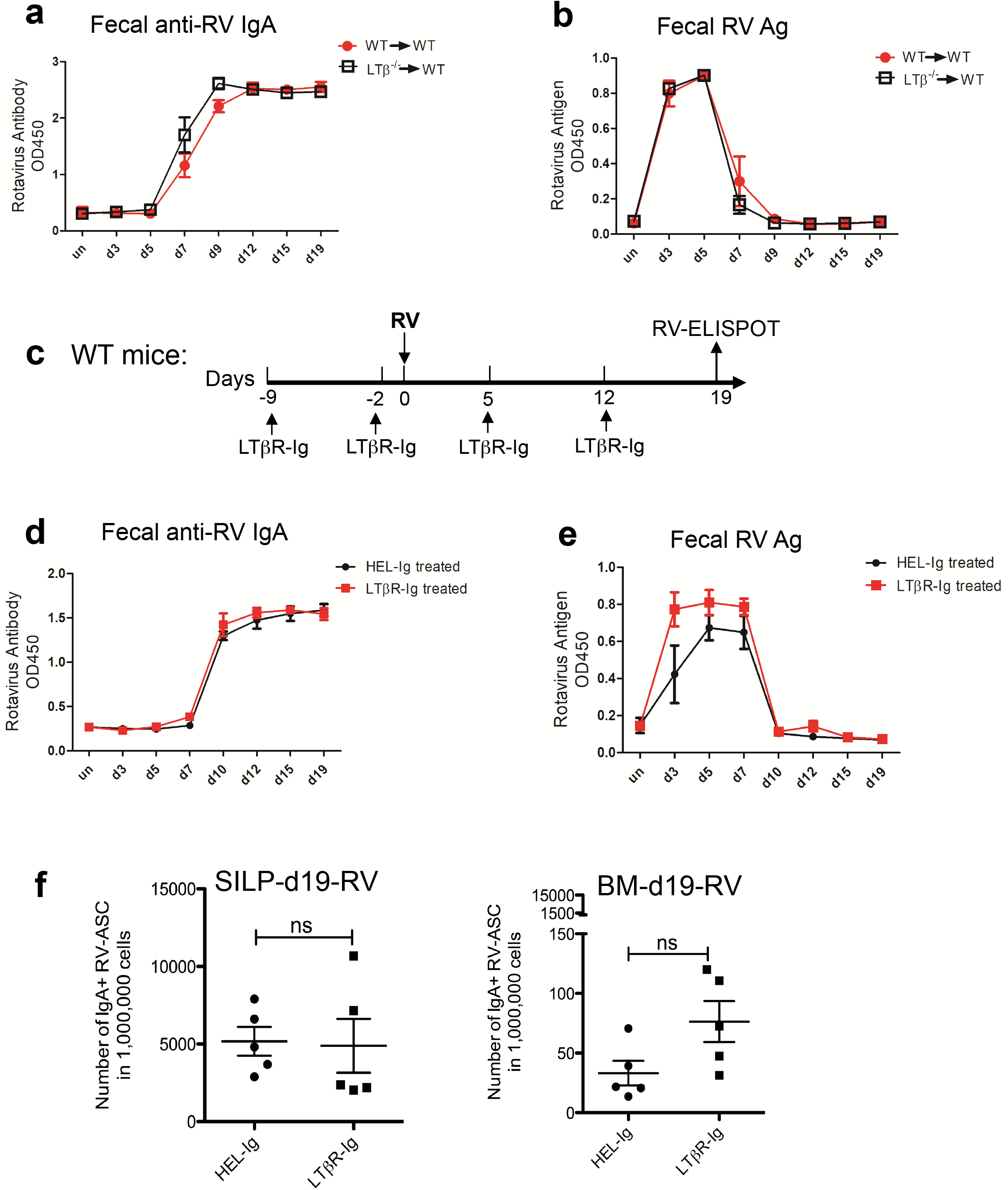
LTβR signalling during adulthood is dispensable for the induction and maintenance of an IgA response against RV. **(a-b)** Representative kinetics of anti-RV IgA induction **(a)** and the presence of RV Ag **(b)** in fecal pellets of the indicated BM chimeric mice as detected by ELISA. Data represent two independent experiments each with three to four mice per time point. All the fecal samples in **(a)** or **(b)** were tested in one ELISA plate. **(c)** Schematic outline of adulthood LTβR-Ig treatment before and during RV infection. **(d)** Levels of fecal anti-RV IgA titres over time in adult LTβR-Ig or control treated mice. **(e)** Levels of fecal RV Ag over time in LTβR-Ig or control treated mice. **(f)** Number of RV-specific IgA^+^ ASC in the SILP and BM of adulthood LTβR-Ig or control treated mice at d19 post-infection. Data in **(c-f)** represent two independent experiments each with four to five mice per group.

**Supplementary Figure 4.**
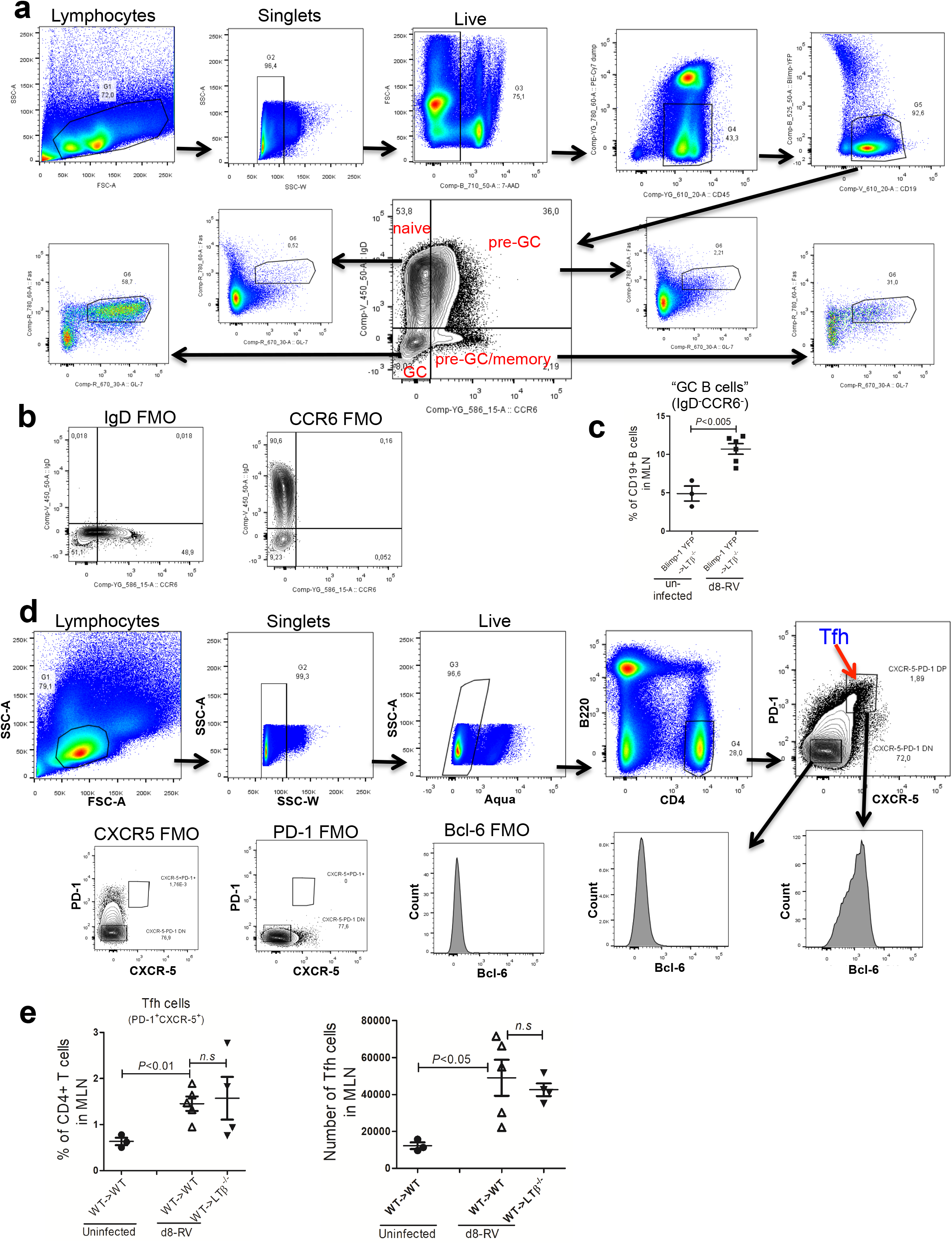
Analysis of germinal center (GC) B cells and follicular T helper (Tfh) cells in the MLN. **(a)** Gating strategy for identification of germinal center B cells in the MLN at d8 post-infection. GC B cells were classified as IgD^−^CCR6^−^, while pre-GC/memory B cells were classified as IgD^−^/CCR6^+^. **(b)** FMO-derived background staining for IgD and CCR6. **(c)** Analysis of the relative percentage of GC B cells in total MLN B cells from uninfected and d8-RV infected WT➔LTβ^-/-^ chimeric mice. **(d)** Gating strategy for the identification of Tfh (CD4^+^/B220^−^CXCR5^+^/PD-1^+^/i.c. Bcl-6^+^) in MLN. i.c.: intracellular. FMO-derived background staining for CXCR5, PD-1 and Bcl-6 are shown. **(e)** Frequency (left panel) and numbers (right panel) of CD4^+^/B220^−^CXCR5^+^/PD-1^+^ Tfh in the MLN of WT➔WT and WT➔LTβ^-/-^ chimeric mice. Data are representative of two independent experiments with 3-5 mice per group.

**Supplementary Figure 5.**
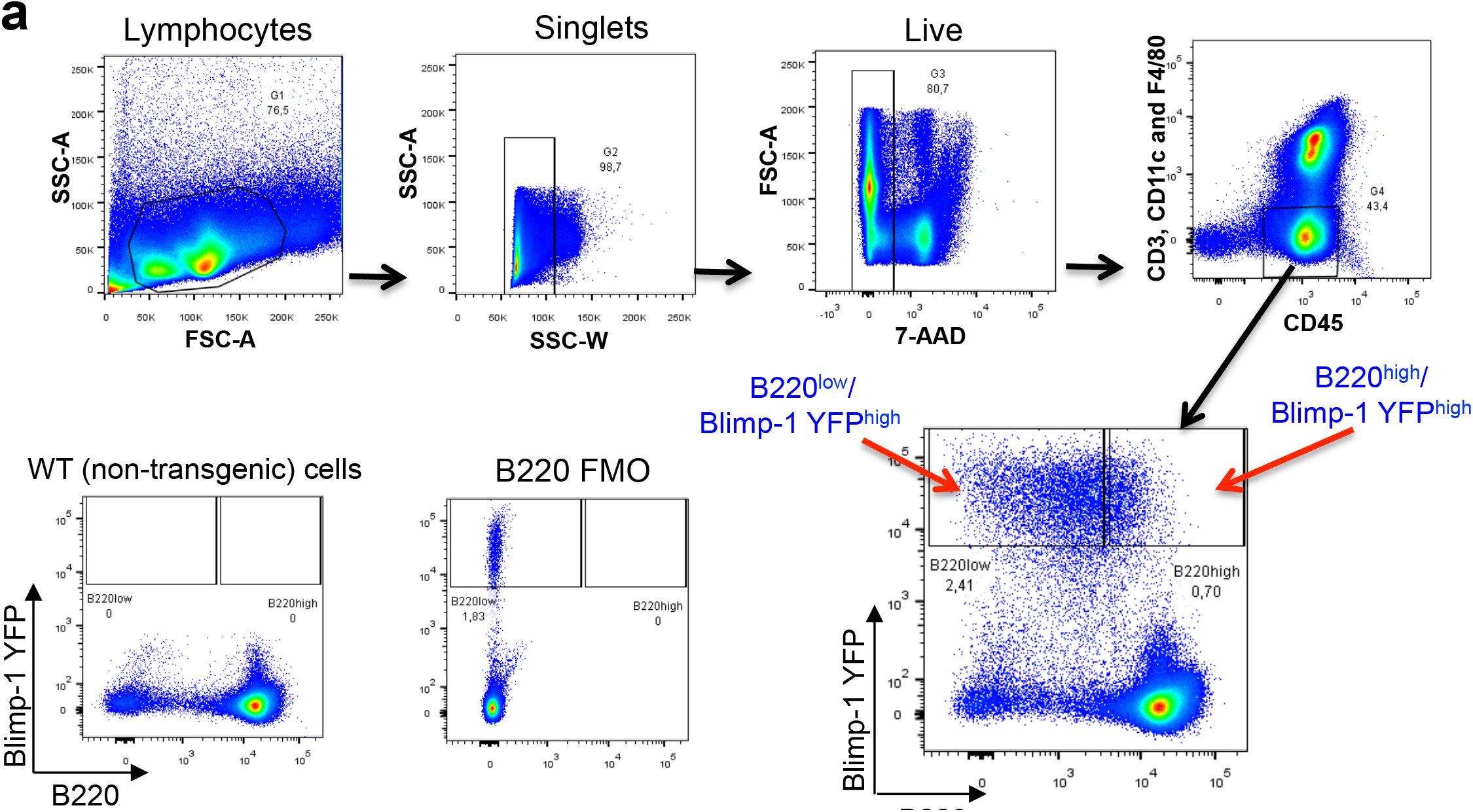
MLN plasma cell analysis post-RV infection. **(a)** Gating strategy for identification of YFP^high^ B cells in the MLN of Blimp-1^YFP^➔WT chimeric mice. Two subsets of YFP^high^ B cells were noted in the MLN at d8 post-infection: CD45^+^/CD3^−^CD3^−^/F4/80^−^/B220^low^/YFP^high^ and CD45^+^/CD3^−^/CD11c^−^/F4/80^−^/B220^higl^/YFP^high^. Background YFP fluorescence (using WT cells) and FMO-derived background staining for B220 are shown.

**Supplementary Figure 6.**
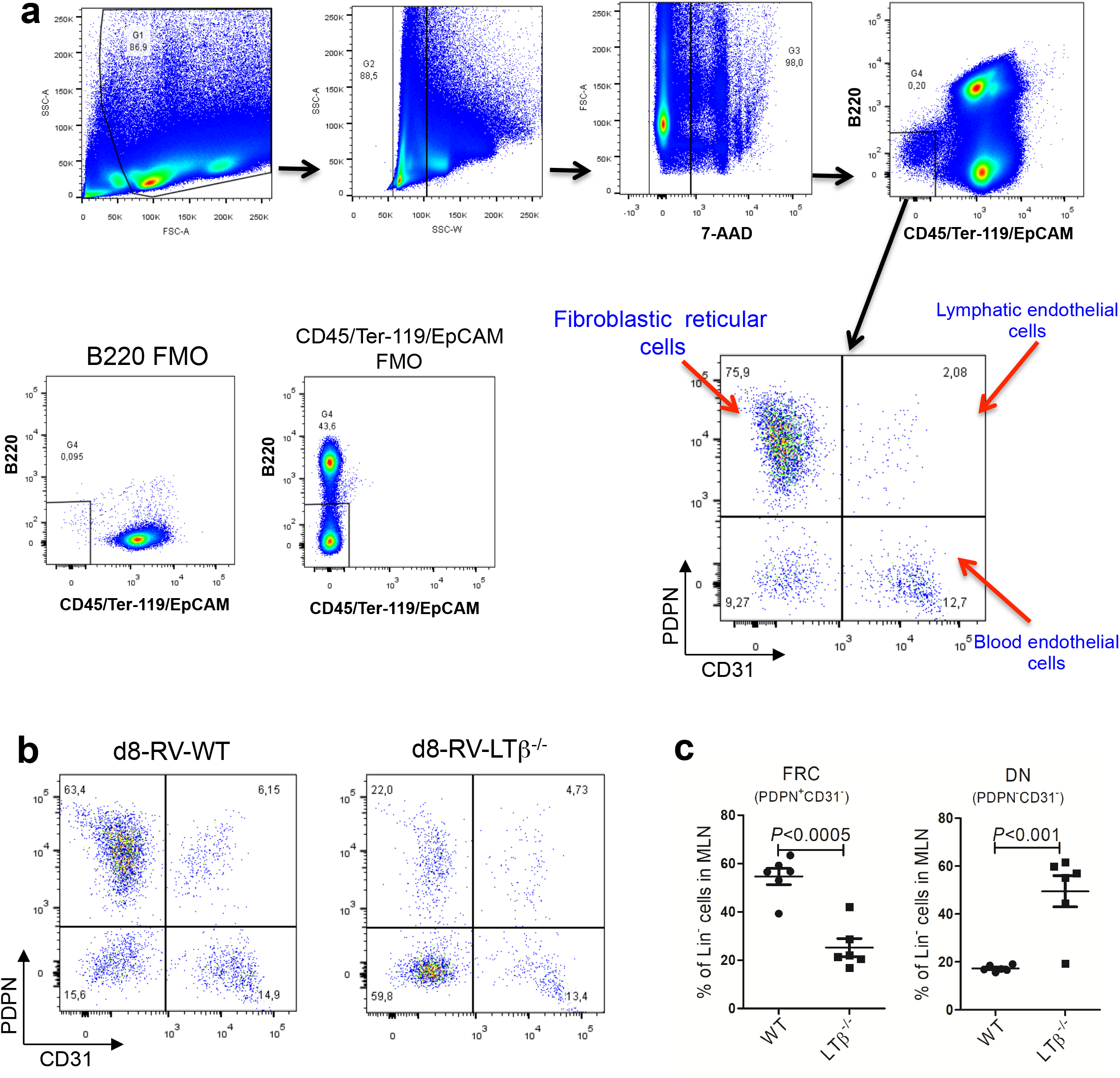
Analysis of lymphoid stromal cells in the MLN. **(a)** Gating strategy for MLN-resident stromal cells. Fibroblastic reticular stromal cells (FRC) were identified as CD45^−^Ter-119^−^/EpCAM^−^/B220^−^/PDPN^+^/CD3^−^, and double negative (DN) stromal cells were identified as CD45^−^/Ter-119^−^/EpCAM^−^/B220^−^/PDPN^−^/CD3^−^. FMO staining of B220 and CD45/Ter-119/EpCAM are shown. **(b)** Representative FACS plots of MLN stromal cells from d8-RV infected WT and LTβ^-/-^ mice. **(c)** Frequency of FRC and PDPN/CD31 DN stromal cells among lineage-negative cells in MLN from WT and LTβ^-/-^ mice at d8 post-infection.

**Supplementary Figure 7.**
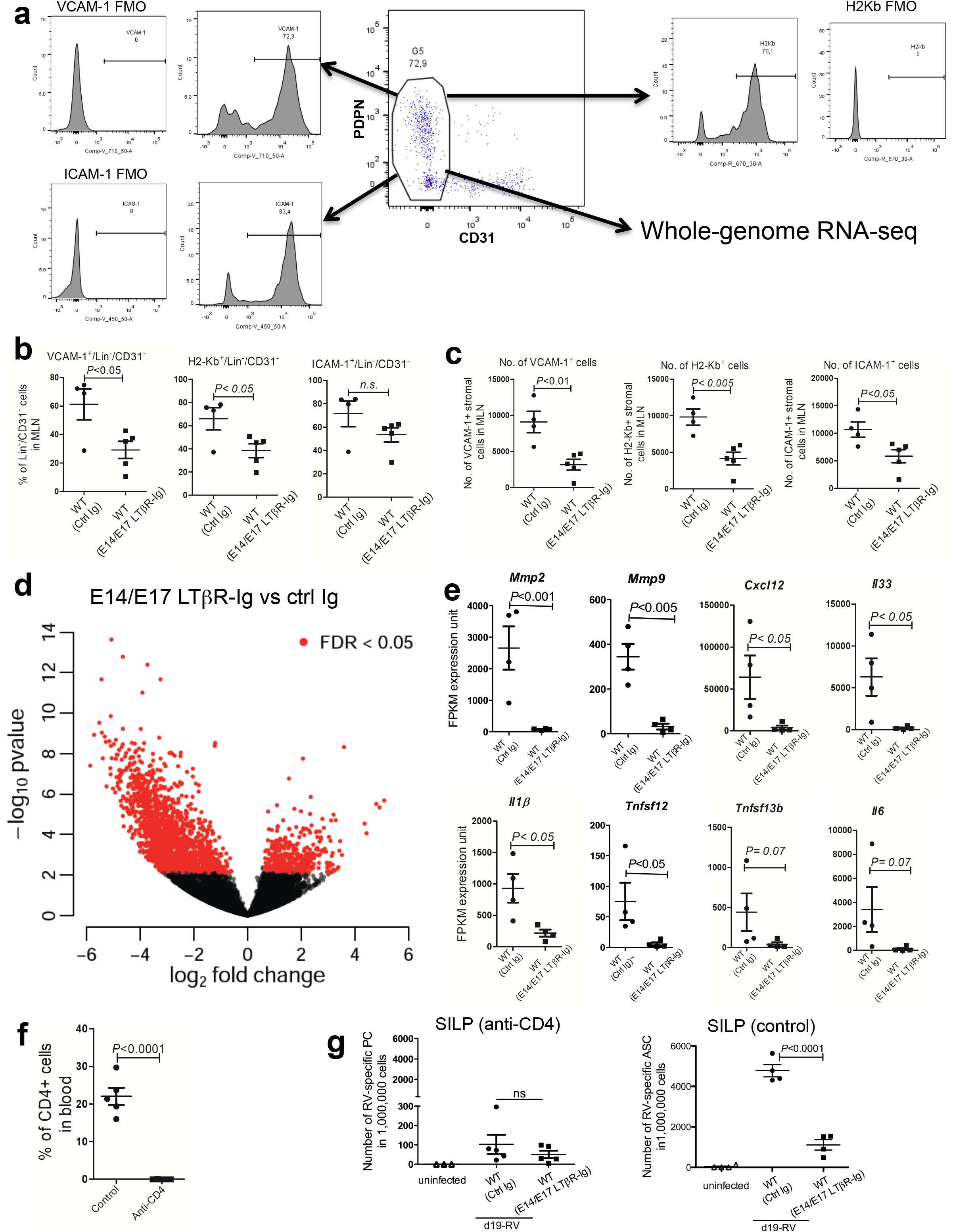
Activation status and gene expression of MLN stromal cells, and the effects of CD4^+^ T cell depletion on the anti-RV IgA response. **(a)** Gating strategy used to assess MLN stromal cell activation markers. CD45^−^/Ter-119^−^/EpCAM^−^/CD19^−^/CD3^−^ stromal cells were gated for analysis of the frequency of VCAM-1, H2-Kb (MHC-I), and ICAM-1, positive cells, respectively. (b-c) Analysis of the frequency **(b)** and numbers **(c)** of VCAM-1^+^, H2-Kb^+^ and ICAM-1^+^ stromal cells among Lin^−^CD3^−^ MLN cells at d8 post-infection of WT mice that received LTβR-Ig or control Ig at E14/E17. **(d)** Whole-genome RNA-sequencing of CD45^−^Ter-119^−^/EpCAM^−^/CD19^−^/CD31^−^ stromal cells sorted from MLN of d8 post-infection WT mice that were treated at E14/E17 with LTβR-Ig or control Ig at. Points colored in red denote genes that were significantly differentially expressed (FDR < 0.05 and fold change > 1.5). The differential expression analysis was performed using DESeq2 (n= 4 samples per group, and each sample is pooled from 2-4 mice). **(e)** RNA-sequencing data for *Mmp2, Mmp9, Cxcl12, Il33, Tnfsfl3b, Il1b, Tnfsf12* and *Il6* from MLN stromal cells derived from mice treated with LTβR-Ig *versus* control Ig at E14/E17. **(f)** Frequency of CD4^+^ T cells derived from peripheral blood lymphocytes from mice that had or had not received anti-CD4 treatment. n=4-5 mice per group. **(g)** Frequency of IgA^+^ RV-ASC in the SILP of WT mice that received LTβR-Ig or control Ig at E14/E17 at d19 post-infection. The left panel shows mice that received anti-CD4 depletion antibody before and during RV infection, while the right panel are mice that did not receive anti-CD4 depletion antibody. Data represent at least two independent experiments each with three to five mice per group.

